# Within-generation and transgenerational social plasticity interact during rapid adaptive evolution

**DOI:** 10.1101/2021.10.21.464843

**Authors:** Samantha L. Sturiale, Nathan W. Bailey

## Abstract

1. The role of within-generation phenotypic plasticity (WGP) versus transgenerational plasticity (TGP) during evolutionary adaptation are not well understood, particularly for socially-cued TGP.
2. We tested how genetics, WGP, and TGP jointly influence expression of fitness traits facilitating adaptive evolution in the field cricket *Teleogryllus oceanicus*. A male-silencing mutation (“*flatwing”*) spread to fixation in ca. 50 generations in a Hawaiian cricket population attacked by acoustically-orienting parasitoids. This rapid loss of song caused the social environment to dramatically change.
3. Juveniles carrying the *flatwing* (*fw*) genotype exhibited greater locomotive activity than those carrying the *normal-wing* (*nw*) allele, consistent with genetic coupling of increased locomotion with *fw*.
4. Consistent with adaptive WGP, homozygous *fw* females developing in the absence of song showed reduced body condition and reproductive investment at adulthood.
5. Adult but not juvenile offspring exhibited TGP in response to maternal social environment for structural size, somatic condition, and reproductive investment, whereas adult locomotion and flight was only influenced by WGP. WGP and TGP interacted to shape multiple traits at adulthood, though effect sizes were modest.
6. Interactions between genetic effects and social plasticity within and across generations are likely to have influenced the evolutionary spread of flatwing crickets. However, interactions among these effects can be complex, and it is notable that TGP manifested most strongly later in development. Our findings stress the importance of evaluating trait plasticity at different developmental stages and across generations when studying phenotypic plasticity’s role in evolution.

## Introduction

Dissecting how phenotypic plasticity affects trait expression within and across generations is necessary to fully understand its role in adaptive evolution. Within-generation plasticity (WGP), where an individual’s phenotype shifts as a response to its own environmental conditions, has long been argued to influence evolutionary processes (Robinson and Dukas 1999; Huey et al. 2003; West-Eberhard 2003; Ghalambor et al. 2007; Lande 2009; Chevin et al. 2010). More recently, researchers have explored plastic responses to parental or grandparental environmental conditions through transgenerational plasticity (TGP) (LaMontagne and McCauley 2001; Dyer et al. 2010; Sheriff et al. 2010). It is clear from this work that TGP can also modify a species’ evolutionary trajectory. In particular, both WGP and TGP may mitigate indirect fitness costs caused by new genetic variants under selection (negative pleiotropy), thus facilitating spread and fixation of *de novo* adaptive variants. However, few empirical studies of rapid adaptation have considered the consequences of both these forms of plasticity acting simultaneously.

Depending on the nature of the interaction between WGP and TGP, various evolutionary outcomes could occur. For example, a study of anti-predator defensive helmet formation in *Daphnia cucullate* found that WGP and TGP additively contribute to offspring phenotype (Agrawal et al. 1999). Such a relationship could move a population towards a new trait optimum faster than if just one form of plasticity were acting (Auge et al. 2017). Several other studies have detected a more complicated pattern, where parental environment interacts with the effects of offspring environment (i.e., TGP alters the extent and/or direction of offspring WGP response or reaction norm) (Prasad et al. 2003; Donelan and Trussell 2015; Luquet and Tariel 2016; Stein et al. 2018; Zirbel et al. 2018). This interaction could be adaptive, for example, by allowing parents to produce pre-adapted offspring that do not need to themselves express costly plasticity (Luquet and Tariel 2016). Alternatively in cases where TGP is non-adaptive for offspring, WGP acting on offspring traits in the opposite direction as TGP could allow offspring a means of ‘escaping’ negative fitness effects carried over from parental environment (Auge et al. 2017). The relationship between TGP and WGP has been somewhat explored in response to the environmental effects of predation (Agrawal et al. 1999; Donelan and Trussell 2015; Luquet and Tariel 2016; Stein et al. 2018), nutrition (Prasad et al. 2003; Zirbel et al. 2018), and temperature (Bernareggi et al. 2016), but the nature of this interaction in other important contexts, such as the social environment, remains largely unknown.

To experimentally dissect this interaction, we tested phenotypic and fitness effects of maternal and offspring social environment in a rapidly-evolving population of the Oceanic field cricket, *Teleogryllus oceanicus*. In Hawaii, acoustically signaling males are attacked by an acoustically orienting parasitoid fly, *Ormia ochracea*. Recently, a single-locus, X-linked mutation, *flatwing* (*fw*), arose and spread in fewer than 20 generations to affect ca. 90% of males in a population on the island of Kauai (Zuk et al. 2006). *Fw* segregates as a single-locus mutation and disrupts normal wing development, thus silencing ‘flatwing’ males and shielding them from fly attack (Zuk et al. 2006; Pascoal et al. 2020). Its rapid spread to fixation dramatically changed the social environment by eliminating the conspicuous long-range male acoustic signal that functions in mate attraction, courtship, and intrasexual aggression. Thus, the mutation has obvious negative indirect (pleiotropic) effects: flatwing males cannot acoustically advertise for mates. They also show partial, apparently maladaptive feminization of phenotypes unrelated to wings (Bailey et al. 2010; Pascoal et al. 2016; Pascoal et al. 2018; Rayner et al. 2019b; Pascoal et al. 2020). Pre-existing WGP to acoustic cues in the environment that offsets some of these fitness costs may have facilitated the rapid spread of *flatwing*. When raised in an environment lacking song, females relax mate preferences and increase responsiveness to calling males, likely enabling them to xslocate the few remaining singing males, or satellite flatwing males near them, in flatwing-dominated populations (Bailey and Zuk 2008; 2012). Similarly, males raised in silence are more likely to express satellite mating tactics (Bailey et al. 2010). Males also show an overall increase in locomotive behavior when reared in silence (Balenger and Zuk 2015). Finally, when exposed to song during rearing, individuals of both sexes develop increased reproductive tissue mass and have enhanced immune responses compared to their counterparts raised in the absence of song (Bailey et al. 2010, 2011; Lierheimer and Tinghitella 2017; Heinen-Kay et al. 2019).

Several features of this field cricket system suggest that socially-induced TGP could also contribute to the rapid spread of flatwing males. Like many insect species, *T. oceanicus* suffers high juvenile mortality, favoring fitness-increasing alterations to early juvenile phenotype via TGP. Recent findings suggest that nongenetic inheritance can affect the interplay between predation risk vs. movement towards singing males, though this has only been examined for adults (Moschilla et al. 2021). Second, the fragmented distribution of *T. oceanicus* habitat in Hawaii, the inability of juveniles to fly, and overlapping generations should result in high autocorrelation between parental and offspring social environments, which is predicted to favor adaptive TGP (Leimar and McNamara 2015). Third, crickets do not possess a fully-developed auditory system until adulthood, though late juveniles may have limited auditory capabilities (Young and Ball 1974; Yack 2004; Staudacher 2009). Young juveniles are therefore not likely capable of accurately assessing their own social environment acoustically, which theory predicts will favor the evolution of TGP (Leimar and McNamara 2015).

We performed three experiments to dissect the potential contributions and interactions of genetic evolutionary responses, WGP, and TGP to rapid adaptation observed in Hawaiian *T. oceanicus*. Supplementary figure S1 provides an experimental overview. Across these experiments, we focused on four traits relevant to mate competition and the loss of acoustic sexual signaling during the evolutionary spread of flatwing crickets: structural size (in juveniles and adults), body condition (in adults), investment in reproductive tissues (in adults), and locomotive activity (in juveniles and adults). Locomotion was a key trait because there is evidence from other insect species that individuals exposed to crowded conditions increase dispersal themselves (WGP) or produce offspring with increased dispersal tendencies (TGP) (Denno and Roderick 1992; Allen et al. 2008; Yu et al. 2019). Locomotive activity is also expected to have especially important fitness consequences in the cricket system given the challenges of locating conspecifics in a song-less, flatwing-dominated population.

In **Experiment 1**, we measured how juvenile locomotive behavior varied across *fw* and *normal-wing* (*nw*) genotypes, before any maternal social manipulation. This allowed us to determine whether the rapid spread of flatwing males might have been associated with genetic changes in juvenile expression of a trait relevant to the changing social environment. We expected that *fw-carriers* benefit from dispersing less as juveniles and therefore aggregating at higher densities upon reproductive maturity, increasing the chances of mating in a song-less environment. In **Experiment 2**, we investigated WGP to the acoustic social environment. We focused on *fw-*carrying individuals because the source population is now all-flatwing (Tinghitella et al. 2018; Rayner et al. 2019a) and previous work indicated that the mutation is associated with increased socially-induced WGP (Pascoal et al. 2018). We tested whether, consistent with previous studies on females carrying *nw* genotypes, *fw*-carrying females (the maternal generation) raised in different acoustic environments increase reproductive investment and alter mating behaviors in a way that increases probability of mating with a silent male. Finally, in **Experiment 3**, we tested transgenerational consequences of the maternal social environment by measuring size and locomotive activity in acoustically-naïve juveniles, and final size, somatic condition, reproductive investment, and locomotive activity in adult offspring. In this last experiment, adults were exposed to either matched or mis-matched acoustic cues compared to their mothers, permitting a direct test of the interaction between WGP and TGP.

## Methods

### Experiment 1: Genotypic differences in juvenile locomotion

#### Cricket populations and rearing

We compared early juvenile behavior across the two cricket morph genotypes using 6 laboratory stock lines – 3 pure-breeding for *fw* and 3 pure-breeding for *nw*. Lines were established in 2016 from a series of controlled crosses of Kauai-derived individuals to ensure homozygosity (Pascoal et al. 2016). Stock crickets were kept in 16 litre plastic containers with cardboard egg cartons for shelter. Twice weekly, they were provided *ad libitum* food (Burgess Supa Rabbit Exel Junior pellets; blended for juveniles) and moistened cotton for water and oviposition. Crickets in isolated-rearing conditions were kept in 100 mL plastic deli pots with shelter, food, and water as above. All subjects were kept in the same growth chamber at 25°C on a photo-reversed 12:12 hour light:dark cycle unless otherwise indicated. To obtain juveniles for this experiment, we collected eggs from each line twice weekly for 4 weeks. After approximately two weeks we monitored egg pads daily (16:00 - 18:00) and isolated new hatchlings.

#### Open-field test

An open field test (OFT) was used to track individual crickets’ movements in an unobstructed arena and measure their total distance travelled, a useful proxy for measuring behaviors related to dispersal, mate location and foraging (Fraser et al. 2001; Dingemanse et al. 2003; Korsten et al. 2013). For this experiment, juveniles were isolated at hatching, and each was tested in an OFT at 15-days and 45 days post-hatching. Juveniles of these ages do not have mature hearing structures (Young and Ball 1974). All OFTs were performed under red light during the dark portion of the crickets’ 12:12 light:dark cycle, between 23-25°C. Subjects were placed in small glass vials within their deli pot to reduce handling disturbance before testing. The vial was gently turned over onto the center of an 11×17 cm clear plastic arena atop white poster paper and the cricket was allowed to acclimatize for two minutes. Upon lifting the vial, we began recording for 5 minutes at 30 frames/second using a camera (Nikon D3300) mounted ca. 40 cm above the arena. The arena was wiped down with 70% ethanol before each trial to minimize residual chemical cues. Two crickets were assayed at once in side-by-side arenas. It is unlikely that they were aware of one another due to their inability to see in red wavelengths of light. After the OFT, each cricket was photographed overtop a micrometer using a Leica DFC295 digital camera affixed to a Leica M60 dissecting microscope. ImageJ (v.1.8.0_112) was used to record pronotum length (a proxy for structural size) from the images.

#### Locomotion measurements

We used DORIS v.0.0.17 (Friard 2019) to extract coordinates of the test subject within each video frame, followed by coordinate path smoothing implemented in R (R Core Team 2020) to increase measurement precision (see Supplemental Methods and Figure S2). Using these coordinates, we measured total distance traveled (“*distance*”) during trials. In this and later experiments involving open field tests, we also explored other movement parameters (“*proportion explored*” as a measure of exploratory activity; and “*origin time*”, “*middle time*”, and “*edge time*” as measures of space usage and thigmotaxis). However, variation in these parameters was largely accounted for by overall differences in distance moved, confirming that *distance* was the most salient locomotion trait in the experiment. For completeness, we discuss the measurement and analysis of all other movement traits in the Online Supplementary Information.

#### Statistical analyzes

All statistical tests were carried out using R version 4.0.2 (R Core Team 2020). We compared *distance* between wing morph genotypes in 15-day old and 45-day old offspring using a linear model. Individuals who jumped during their assay (n = 2 in 45-day assay) or whose video was inadvertently deleted before analysis (n = 2 in 45-day assay) were excluded. All data transformations are shown in Table S1.

Morph and sex were modelled as categorical variables, with line nested within morph to account for inter-line variation. Pronotum length, temperature, and time of day were included as covariates. Thirty-nine individuals died before their sex could be identified, so to verify that sex did not qualitatively affect the findings, models including sex as a fixed effect were run on the subset of individuals for which sex could be identified. Sex did not approach significance in this model (all *p* > 0.2) and the qualitative outcome did not differ. Thus, the model retaining all individuals, and excluding sex as a fixed effect, was retained (Equation 1 of Supplementary Table S2). Finally, the model was run first with all individuals, then with only those who moved during the assay to confirm that genotypic variation in *distance* was not due to differences in the likelihood of initiating movement. Excluding crickets that failed to initiate movement did not affect interpretations of genotype differences, so final models included stationary crickets (Supplementary Table S3).

### Experiment 2: Social plasticity in the maternal generation (WGP)

#### Cricket populations and rearing

The three pure-breeding *fw* lines used in Experiment 1 were reciprocally interbred to create an admixed pure-breeding *fw* stock population. Following previous work, we isolated juvenile females from this stock when sex became apparent to ensure virginity and more easily manipulate their acoustic environment (Bailey and Zuk 2008; Pascoal et al. 2018). We also segregated a group of juvenile males into single-sex 16-L box to maintain their virginity. All group rearing conditions were identical to Experiment 1. Isolated females were placed in a separate, temperature-controlled 25°C incubator on a 12h:12h photo-reversed light cycle, with no male calling. Females were checked daily for adult eclosion, whereupon they were haphazardly assigned one of two acoustic social treatments: Song or No Song. Females do not achieve reproductive maturity until several days after adult eclosion, so our acoustic treatment targets the developmental period when mate assessment is possible but mating is not (Swanger and Zuk 2015). We also recorded the number of days spent isolated prior to eclosion to account for any differences in growth rate that might be associated with time spent without song prior to adult acoustic treatment. We kept each female in their acoustic treatment for 15 days post-eclosion.

#### Acoustic treatments

In the Song treatment, Kauai male calls reflecting population averages for key song parameters were played at 80-85 dB (measured at the lid of the deli cup which has an acoustic impedance of ca. 10 dB) during the night portion of the crickets’ light:dark cycle to best match calling dynamics in the wild (Zuk et al. 1993). Playbacks used in the Song treatment have been previously described (Pascoal et al. 2018) (see Supplementary Methods). Acoustic treatments were run in two separate LMS Series 4 (Model 600) controlled temperature incubators at 25°C on the same 12h:12h photo-reversed light:dark cycle as the general incubator. Calls were broadcast from computer speakers (Logitech Z120 2.0) and the calling schedule programmed using the Task Scheduler application on a desktop computer. Twice a week, we switched which incubator housed each acoustic treatment to prevent any incubator-related experimental confounds.

#### Mating trials of acoustically treated females

At 15 days post-eclosion, isolated adult females were weighed and their pronotum width was measured using digital calipers. Each female was placed in a 16 × 18 cm plastic container with cardboard, rabbit chow, and moistened cotton. We haphazardly selected an adult virgin male from the flatwing stock population, weighed it, measured its pronotum width, and placed it in the container with the female. Trials were performed between 20-23°C under red light between 16:00h and 18:00h. They lasted for 20 minutes, and we noted whether the female mounted the male and whether the male transferred a spermatophore. Afterwards, pairs were placed in a separate incubator without male song at the same temperature and light:dark cycle as in Experiment 1. After 24 hours, the male was removed to reduce potential paternal influences on offspring phenotype. After another 24-48 hours, the female was removed, and the egg pad was collected for use in Experiment 3.

#### Body condition, size, and reproductive tissue measurements

To compare female body condition, we used pronotum width and total body weight to calculate the scaled mass index (SMI) of each individual (Peig and Green 2009). A subset of females drawn haphazardly from each treatment (total n = 23) were dissected at 15 days post-eclosion rather than mated. We recorded their pronotum width using digital calipers, weighed them, and then determined wet mass of their dissected ovaries. Somatic mass was calculated by subtracting ovary mass from total weight.

#### Statistical analyzes

First, we compared *SMI* across acoustic treatments using a linear model (Equation 2 of Table S2) with acoustic treatment as a categorical factor, days isolated before treatment as a covariate, and experimental replicate (block one or block two of the experiment). Replicate only had two factor levels so we included it as a fixed effect. Second, we compared mating behavior across acoustic treatments. We first ran a generalized linear model (GLM) with binomial error to examine presence vs. absence of *female mounting* during trials, including acoustic treatment, female SMI, and male SMI as predictors. Next, we ran a binomial GLM examining presence vs. absence of *spermatophore transfer*. For this, we only included the 49 mating trials (out of 65 total) where mounting had occurred, because spermatophore transfer cannot occur without mounting. Acoustic treatment was included as a categorical factor and female SMI and male SMI were included as covariates. Equation 3 in Table S2 gives the general form of these models.

Finally, we compared female reproductive investment (ovary mass) across acoustic treatments in the subset (n = 23) of females that had been dissected by running a linear model (LM) on *ovary mass*, with acoustic treatment and days isolated as predictor and covariate, respectively. As in previous studies of reproductive investment, we controlled for body size by including log-transformed soma mass as an additional covariate (Tomkins and Simmons 2002; Bailey et al. 2010) (Equation 4 in Table S2).

### Experiment 3: Transgenerational effects of maternal social environment and interactions between TGP and WGP in adult offspring

#### Cricket populations and rearing

Eggs produced by the maternal generation in Experiment 2 were first kept in a separate incubator under the same temperature and light conditions as the general incubator. As they began to hatch, the first UK national lockdown in response to the 2020 Covid-19 pandemic (23 March 2020) required that all laboratory experiments be run under strict social distancing measures, which affected where and how some of our procedures were executed. A description of ‘socially-distanced’ methods plus our design to statistically account for any variation it introduced is provided in the Supplementary Methods.

We first tested TGP effects in juvenile offspring. Hatchlings were isolated as described previously and kept at 18-24°C on a 12h:12h photo-reversed light:dark cycle. We ran the experiment in two blocks, with individuals in the second block kept at 25°C in the lab incubator, as in Experiment 1. For the early juvenile offspring TGP experiment, we tested 311 offspring (131 from 14 mothers treated with Song, 180 from 21 mothers treated with No Song) at 15 days post-hatching. 199 individuals were tested again at 45 days post-hatching (117 from 11 No Song mothers and 82 from 8 Song mothers).

For adult offspring experiments, hatchlings were kept in 10 replicate group-rearing boxes during development to match the demographic rearing conditions experienced by the previous generation. Once sex was apparent during development, individuals were isolated and assigned to either acoustic treatment using the same incubators and playback schedules as in Experiment 2. The distribution of adult offspring (n = 387) across the 4 maternal-offspring acoustic treatment combinations is shown in Supplementary Table S4. It is important to note that the adults measured in this experiment were not the same individuals as those used for the TGP juvenile trials described above; thus, the corresponding data sets are derived from different individuals of the same generation.

#### Open field test

OFT procedures were identical to Experiment 1. For juvenile offspring, OFTs were performed 15 days post-hatching and 45 days post-hatching. Adult offspring OFTs were performed 8-days post-adult eclosion. All recordings were performed at 23-28°C between 12:00 and 17:00 under dim red lighting. Adult OFTs were identical to those of juveniles, except a larger plastic arena was used (41 cm wide, 37 cm long, and 28 cm high). In the course of the experiment, we noticed that some adults attempted to fly out of the arena during the assay. When that happened, we stopped the recording and placed the subject into an incubator without song for 10 minutes. After re-acclimation, we started the trial again. The number of flight attempts was recorded for each individual. We collected movement coordinates and calculated *distance* using DORIS (v.0.0.17) as in Experiment 1.

#### Morphological measurements

Following the OFT, we photographed each juvenile overtop a micrometer using the same camera and dissecting scope as in Experiment 1 and measured pronotum length using ImageJ (v.1.8.0_112). We euthanized adults after OFTs at 8 days post-eclosion, then weighed them and measured pronotum length to the nearest 0.01 mm using digital calipers. We then dissected, blotted excess fluid, and weighed their gonads (male testes and accessory glands, female ovaries). Here we used pronotum length and soma weight to calculate SMI, by subtracting gonad weight from total weight. In this experiment, SMI was thus a measure of somatic body condition, which allowed us to investigate whether differences in maternal or offspring acoustic environments influenced relative investment in somatic tissues while scaling to structural size. SMI was calculated separately for each sex. Gonad weight was later compared directly.

#### Statistical analyzes

First, we tested whether *juvenile offspring size* differed between maternal acoustic treatments by running a linear mixed model (LMM) using pronotum length as the response. Maternal treatment and experimental replicate were included as fixed effects, with maternal ID as a random effect. Experimental replicate was included to account for different rearing temperatures in trial 1 and trial 2 (see Supplementary Methods). The treatment*replicate interaction (*p* > 0.2) was excluded from the final model. Pronotum length of 45-day old offspring was modelled similarly except experimental replicate was not included because we only had 45-day nymph data for trial 1. Models took the general form of Equation 5 in Supplementary Table S2.

We then examined the effect of maternal treatment on juvenile offspring *distance* using an LLM which included maternal treatment and experimental replicate as categorical factors; temperature, time of day, and pronotum length as covariates; and maternal ID as a random effect. The treatment*replicate interaction (*p* > 0.2) was excluded from the final model. A similar model was run for 45-day old offspring, except experimental replicate was not included because we only had 45-day data for trial 1. The general form of the model is given by Equation 6 in Supplementary Table S2.

To investigate TGP and WGP and their interaction, we tested the effect of maternal and offspring acoustic treatments on *adult pronotum length* and *somatic condition (SMI)*. Because there are large sex differences in physiology and the possibility of sex-specific maternal effects, we ran separate models for each sex. Each LMM included maternal and offspring treatments as factors plus their interaction. Non-significant (*p* > 0.2) interactions were removed. We analyzed the effect of acoustic treatments on *adult offspring reproductive investment* using sex-specific models with gonad weight as the response. First we compared unscaled reproductive investment, then we added pronotum length as a covariate to examine whether variation in reproductive investment could be explained by structural size. Finally, we added log-transformed somatic mass to examine whether variation in reproductive investment might be explained by somatic weight. Replicate was included as a random effect. These models took the general form shown in Equation 7 of Supplementary Table S2.

We then tested the impact of TGP and WGP on adult *distance* using separate LMMs for each behavior and sex. All models included maternal treatment, offspring treatment, temperature, time, and somatic SMI, plus replicate as a random effect (Equation 8 in Table S2). Non-significant (p > 0.2) maternal*offspring treatment interactions were removed.

As a *post hoc* analysis of treatment and sex variation in attempted flight behavior, we modelled *flight attempts* using a generalized linear mixed model (GLMM) with a binomial error (1 if flight was attempted and 0 if not) with the predictors: maternal treatment, offspring treatment, sex, maternal treatment*sex, and somatic SMI. Replicate was included as a random effect. Equation 9 of Supplementary Table S2 describes the final model after removing non-significant interaction terms.

## Results

### Experiment 1: Genotypic differences in juvenile locomotion

*Fw*-carrying nymphs moved further than *nw* nymphs, both at 15 days and 45 days post-hatching (Table 1 Figure 1). The effect size of distance differences was considerable. 15-day-old *fw* nymphs, which had a mean length of 3.81 mm, moved an average of ca. 300 mm further than *nw* nymphs. This difference in distance moved is ca. 79x their body length in a relatively short period of 5 minutes. For 45 day-old nymphs, the average movement differential was 132.73 mm (ca. 12.9x the mean body length of a 45 day-old nymph).

**Table 1.** Linear models examining the effects of genotype on total distance travelled in open field tests by 15 day-old (top) and 45 day-old (bottom) juveniles

|  |  | *df* | *F* | *P*<sup>†</sup> |
| --- | --- | --- | --- | --- |
| 15 days old | Morph | 1, 245 | 12.988 | **<0.001** |
|  | Morph:Line | 4, 245 | 2.806 | **0.026** |
|  | Pronotum | 1, 245 | 0.360 | 0.549 |
|  | Temperature | 1, 245 | 0.680 | 0.410 |
|  | Assay Time | 1, 245 | 0.311 | 0.578 |
| 45 days old | Morph | 1, 210 | 16.554 | **<0.001** |
|  | Morph:Line | 4, 210 | 12.696 | **<0.001** |
|  | Pronotum | 1, 210 | 12.122 | **<0.001** |
|  | Temperature | 1, 210 | 0.561 | 0.455 |
|  | Assay Time | 1, 210 | 33.055 | **<0.001** |

\* numerator, denominator † p < 0.05 indicated in bold

**Figure 1.**
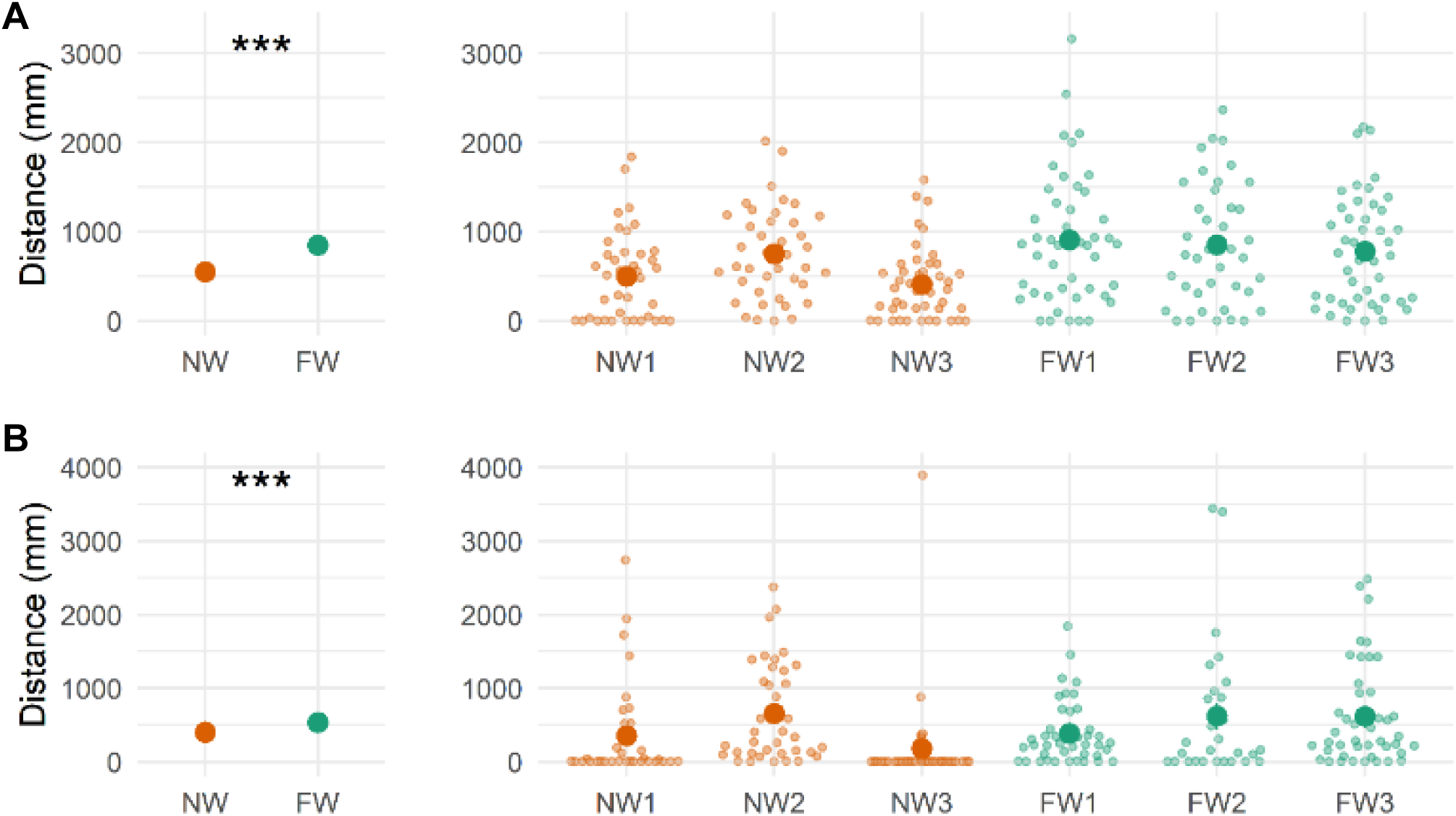
The effect of wing morph genotype on total distance traveled by juveniles in open field tests for (**A**) 15 day-old nymphs, (**B**) 45 day-old nymphs. Plots on the left illustrate pooled means across three replicate morph lines. Bars indicating ± 1 standard error are not shown as these are too small to indicate graphically without being obstructed by the symbols for means. Violin plots on the right show data for each replicate morph line, with dark circles indicating means, small light circles showing each data point, and bars indicating ± 1 standard error (when visible). *** indicates a morph difference with p < 0.001.

### Experiment 2: Social plasticity in the maternal generation (WGP)

The acoustic social environment affected physiology (Table 2 Figure 2), but not mating behavior (Supplementary Table S5; Supplementary Figure S4), in homozygous *fw* females. Those raised in Song attained higher condition (SMI) (Table 2 Figure 2A) and had heavier ovaries relative to somatic mass (Table 2 Figure 2B). Female mounting was not influenced by prior acoustic experience, though females were more likely to mount higher condition males (Supplementary Table S5 and Supplementary Figure S4A). Similarly, in trials where females did mount males (n = 49), there was no evidence that acoustic treatment affected spermatophore transfer, though males were more likely to transfer a spermatophore to higher condition females (Supplementary Table S5 and Supplementary Figure S4B).

**Table 2.** Linear models examining WGP arising from the acoustic environment on female condition (scaled mass index) and reproductive weight (g)

|  | Condition |  |  | Reproductive Weight |  |  |
| --- | --- | --- | --- | --- | --- | --- |
|  | *df*<sup>*</sup> | *F* | *P*<sup>†</sup> | *df* | *F* | *P* |
| Acoustic treatment | 1,110 | 6.038 | **0.016** | 1,19 | 5.015 | **0.037** |
| Days isolated | 1,110 | 0.129 | 0.720 | 1,19 | 1.037 | 0.321 |
| Experimental replicate | 1,110 | 0.948 | 0.332 | - | - | - |
| Somatic mass | - | - | - | 1,19 | 0.211 | 0.652 |

<sup>\*</sup> numerator, denominator
<sup>†</sup> $p < 0.05$ indicated in bold

**Figure 2.**
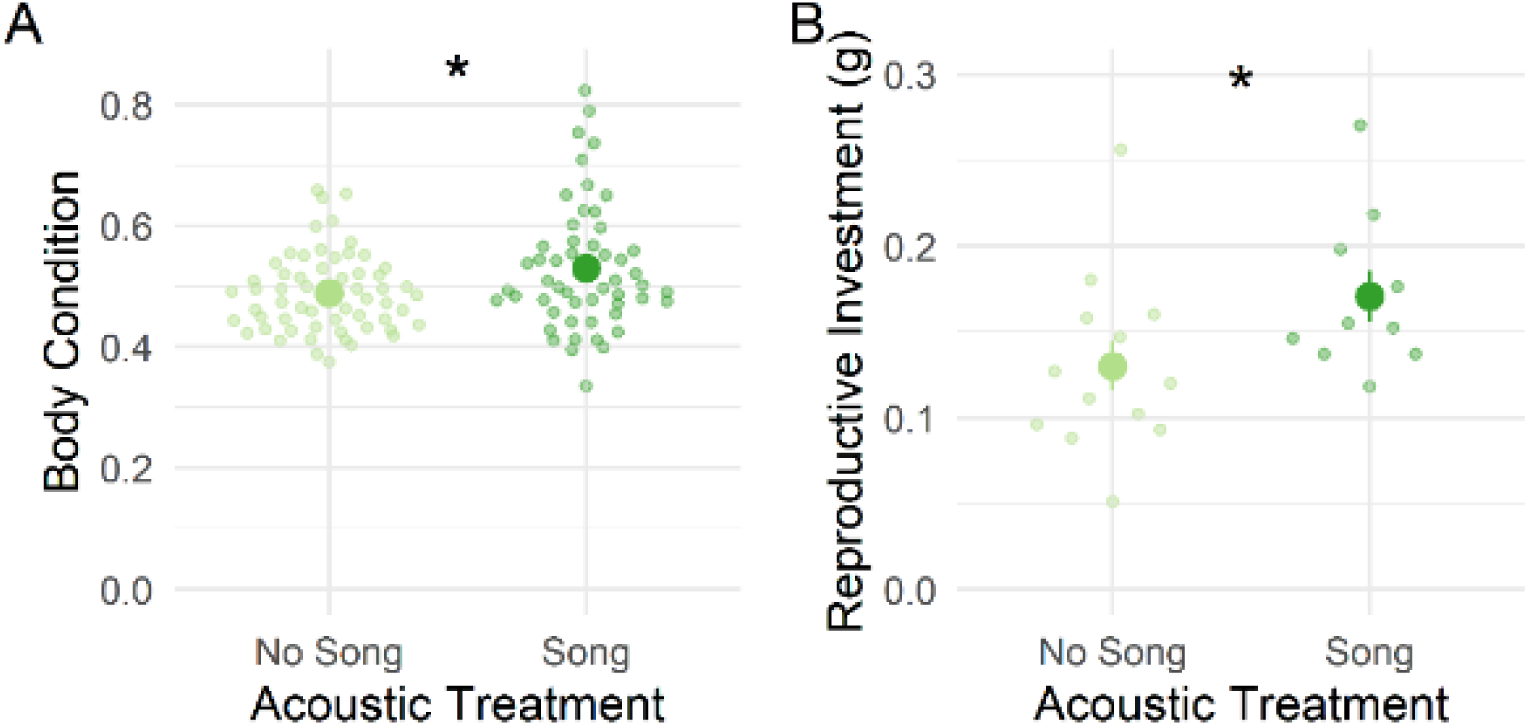
The effect of acoustic environment on (**A**) adult female body condition (scaled mass index) and (**B**) reproductive investment (ovaries mass). Plots show means ± 1 standard error, except for (A), where standard error bars are too small to indicate graphically without being obstructed by the symbols for means. Asterisks indicate p < 0.05.

### Experiment 3: Transgenerational effects of maternal social environment and interactions between TGP and WGP in adult offspring

Unexpectedly, TGP affected adult, but not juvenile, traits (Figure 3). For example, the acoustic treatment of mothers was not associated with locomotion and morphology of their 15-day-old and 45-day old juvenile offspring (Table 4 Supplementary Tables S6 and S7). However, adult offspring that experienced song themselves during rearing moved further (WGP) (Table 5; Figure 3A), and in the case of males, this WGP was considerably exaggerated if their mothers had been raised without song (TGP) (Figure 3A right). It must be noted, this WGP*TGP interaction was only marginally significant (*p* = 0.055; Table 5), though the effect size appears non-trivial (mean movement differential of ca. 400 mm for adult song-reared males; Figure 3A right).

**Table 4.** Linear models examining the effects of TGP arising from the maternal acoustic environment on total distance travelled by 15 day-old (top) and 45 day-old (bottom) juveniles

|  |  | *df* * | *F* | *P*† |
| --- | --- | --- | --- | --- |
| 15 days old | Maternal treatment | 1, 300 | 0.625 | 0.429 |
|  | Experimental replicate | 1, 300 | 45.092 | **<0.001** |
|  | Pronotum | 1, 300 | 0.086 | 0.769 |
|  | Temperature | 1, 300 | 0.530 | 0.467 |
|  | Assay time | 1, 300 | 1.073 | 0.300 |
| 45 days old | Maternal treatment | 1, 191 | 0.003 | 0.955 |
|  | Pronotum length | 1, 191 | 5.714 | **0.017** |
|  | Temperature | 1, 191 | 2.278 | 0.131 |
|  | Assay time | 1, 191 | 0.338 | 0.561 |

\* numerator, denominator
† $p < 0.05$ indicated in bold

**Table 5.**
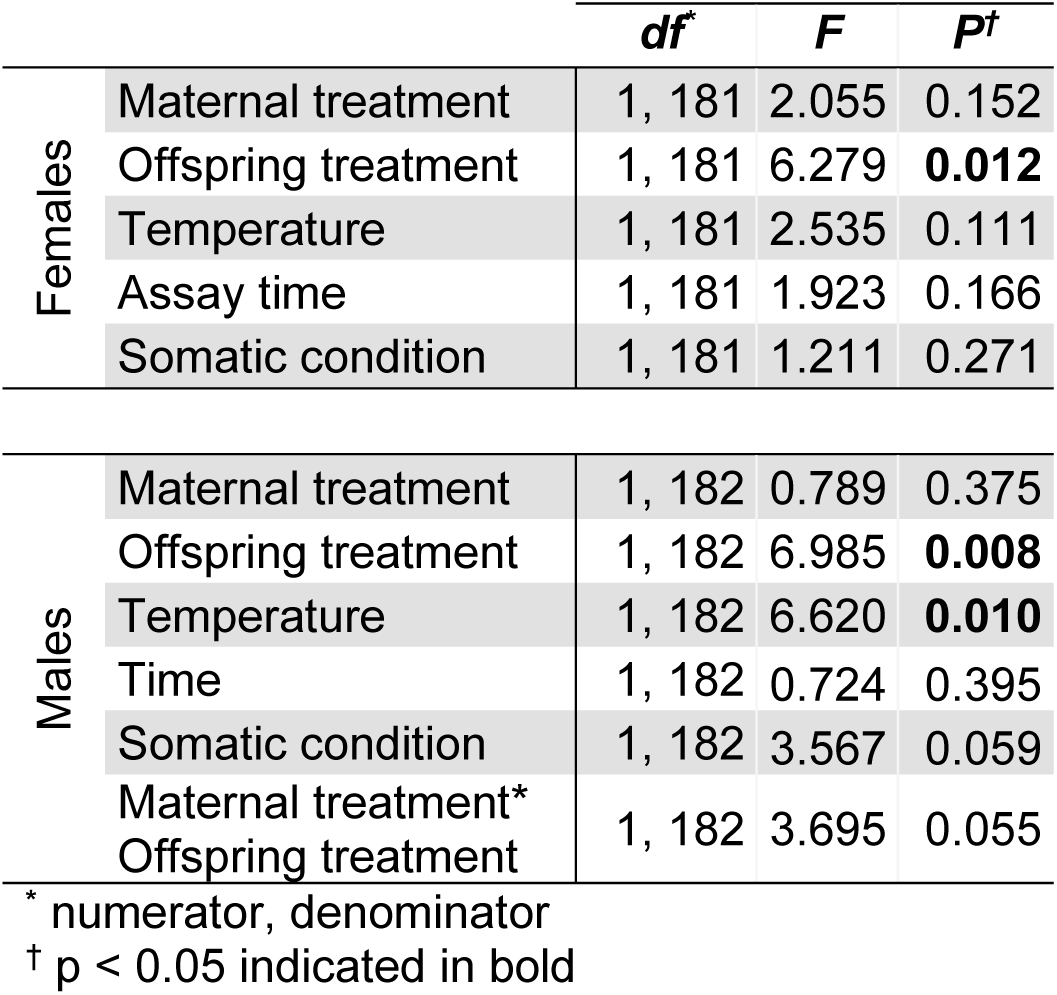
Linear models examining TGP and WGP on total distance moved by adult females (top) and adult males (bottom)

**Figure 3.**
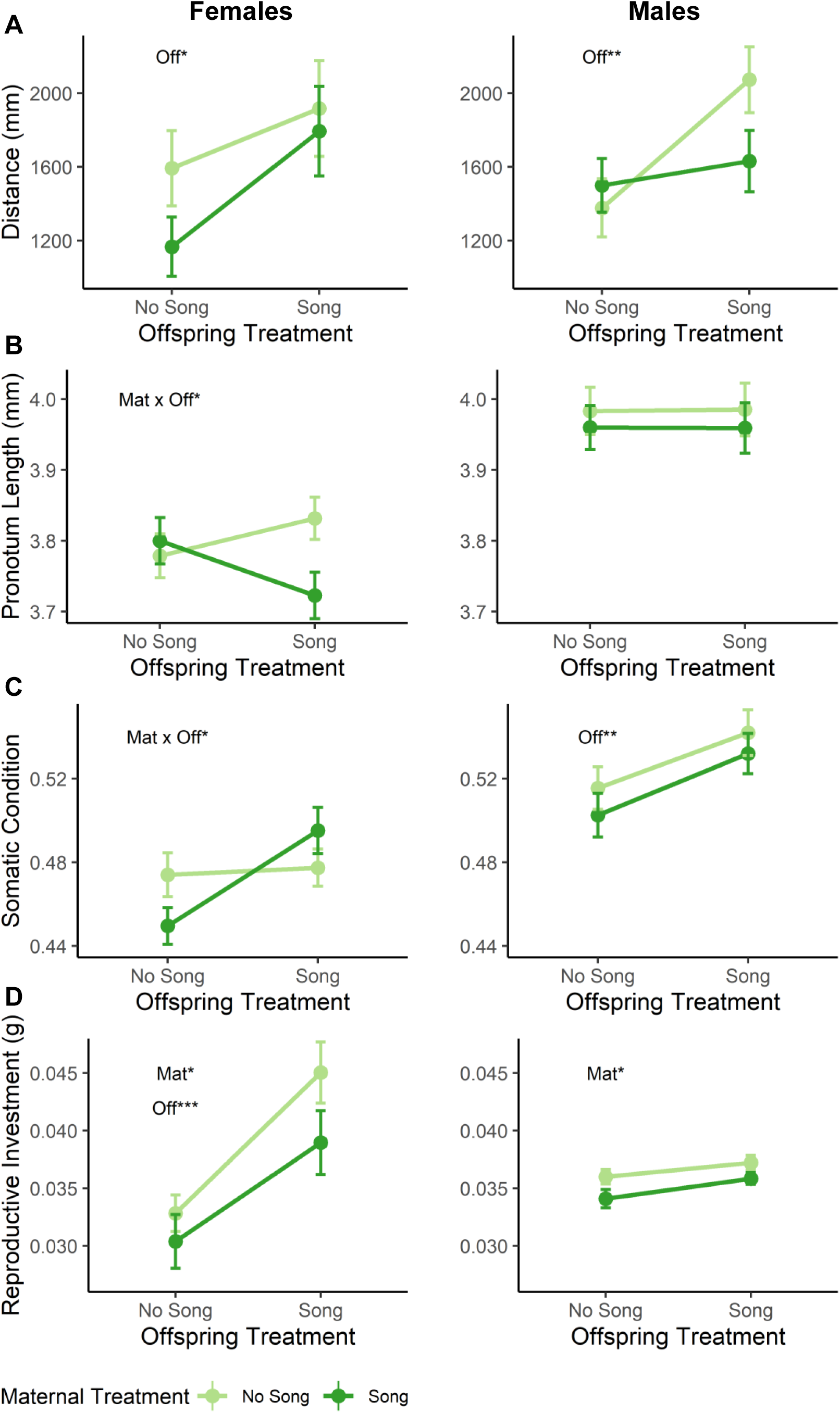
Effects of TGP and WGP in adult female (left) and adult male (right) offspring. (**A**) Distance travelled (**B**) pronotum length, (**C**) somatic condition, (**D**) reproductive investment, i.e., female ovaries mass and male testis mass. Means are indicated by circles and bars indicate ± 1 standard error. Significance of maternal (“Mat”) treatment, offspring (“Off”) treatment, and their interaction (“Mat × Off”) are indicated with asterisks: \**p* <0.05 \*\**p* <0.01 \*\*\**p* <0.001

By contrast, acoustic effects on adult offspring morphology provided strong and consistent evidence for WGP, and TGP also affected several aspects of adult offspring morphology (Tables 6 and 7; Figure 3B,C,D). TGP often did not affect traits in the same direction as WGP; interactions between TGP and WGP combined to shape adult traits in a way that suggests TGP activates WGP, or put another way, that the manifestation of TGP is contingent on current environmental conditions. For example, female offspring reared without song had similar pronotum lengths, but when they were reared with song, those whose mothers experienced No Song grew to be larger than those whose mothers experienced Song (Table 6; Figure 3B left). Female somatic condition showed a similar crossing-over effect (Table 6; Figure 3C left). Also, TGP and WGP affected female investment in ovaries, but in conflicting directions. Those raised in song developed heavier ovaries than those raised without song, and offspring from the No Song maternal treatment developed heavier ovaries than offspring from mothers who experienced Song (Table 7; Figure 3D left). Adult males raised in the presence of song developed higher somatic condition than those raised without song, regardless of maternal treatment (Table 6; Figure 3B right). We also found that crickets raised without song attempted flight more than those raised with song, particularly for females and lower condition individuals (Table 8; Figure 6; Supplementary Figure S5).

**Table 6.**
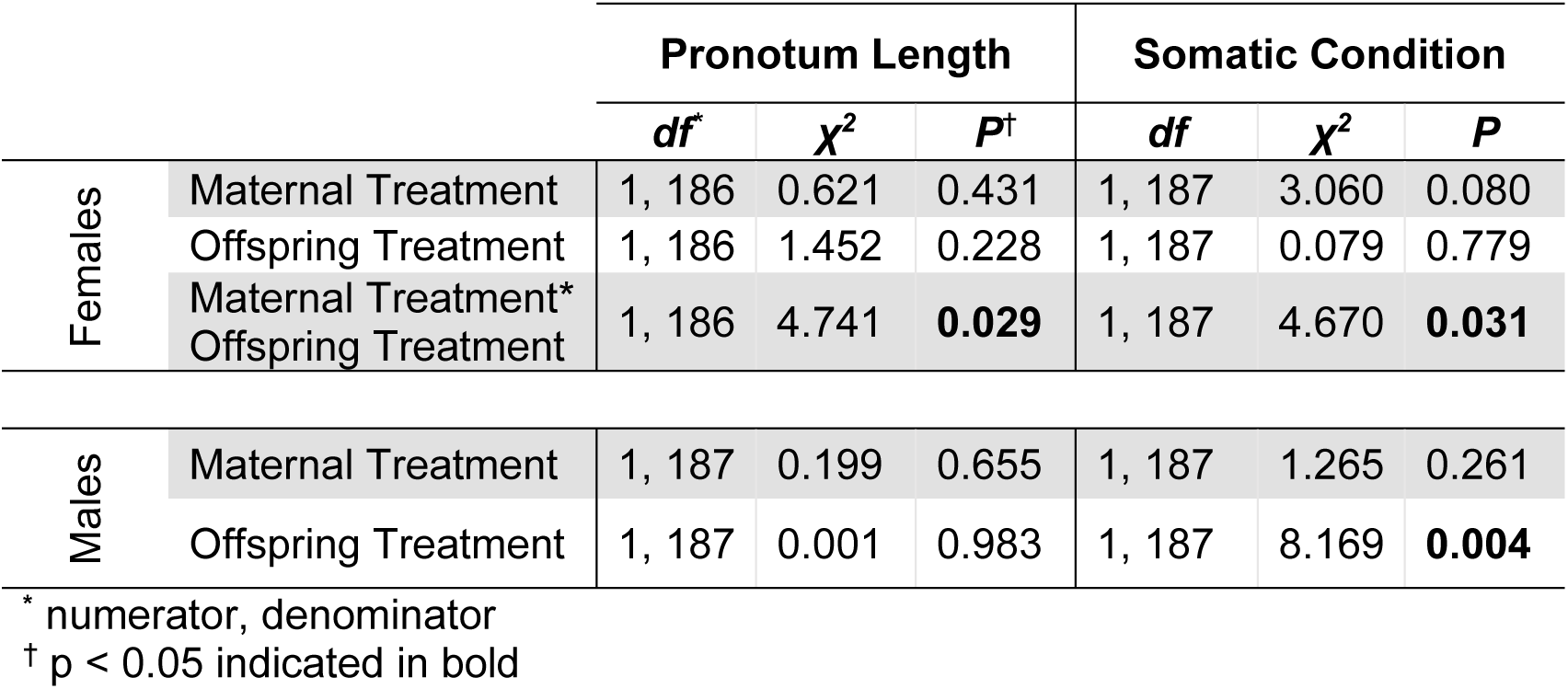
Linear models examining TGP and WGP on adult morphology for females (top) and males (bottom)

**Table 7.** Linear models examining TGP and WGP on reproductive investment for adult females (top) and males (bottom)

| | | *df* | $\chi^2$ | *P* |
| --- | --- | --- | --- | --- |
| Females | Maternal treatment | 1, 186 | 3.910 | **0.048** |
|  | Offspring treatment | 1, 186 | 16.860 | **<0.001** |
|  | Pronotum length | 1, 186 | 0.894 | 0.345 |
|  | Somatic weight | 1, 186 | 15.601 | **<0.001** |
| Males | Maternal treatment | 1, 185 | 3.975 | **0.046** |
|  | Offspring treatment | 1, 185 | 2.800 | 0.094 |
|  | Pronotum length | 1, 185 | 1.698 | 0.193 |
|  | Somatic weight | 1, 185 | 8.675 | **0.003** |

\* numerator, denominator
† *p* < 0.05 indicated in bold

**Table 8.** GLMM examining the effects of WGP and TGP on flight attempts

| | *df* | $\chi^2$ | *P* |
| --- | --- | --- | --- |
| Maternal treatment | 1, 371 | 0.085 | 0.770 |
| Offspring treatment | 1, 371 | 8.788 | **0.003** |
| Sex | 1, 371 | 4.213 | **0.040** |
| Somatic condition | 1, 371 | 4.161 | **0.041** |
| Maternal treatment*Sex | 1, 371 | 2.140 | 0.144 |

\* numerator, denominator
† *p* < 0.05 indicated in bold

## Discussion

Phenotypic plasticity’s role in evolution stimulates vigorous debate, but one barrier to a general resolution may be that plasticity is not a monolithic phenomenon. Influential verbal models have suggested that “buffering” effects of plasticity can permit novel adaptations to escape loss at low frequencies and subsequently spread under selection, but few empirical studies have been able to assess the contributions and interactions of genetics and different forms of phenotypic plasticity such as within-generation and transgenerational plasticity. Here we demonstrate how all three inputs – genetics, WGP and TGP – interact to affect traits that potentially facilitate the rapid evolution of a parasitoid-avoidance adaptation in Hawaiian field crickets, male silence.

Genotype had a surprisingly large effect on juvenile behavior, but in the opposite direction predicted. At very early juvenile stages, *flatwing* carriers moved nearly 80 body lengths further than *normal-wing* carriers in a span of only 5 minutes. This genotypic difference could result from pleiotropic effects of the *fw* mutation or genomic hitchhiking. Genotypic differences in juvenile locomotion are consistent with phenotypic differences that have been detected between *fs* and *nw* carriers in other sexually dimorphic adult traits (Pascoal et al. 2016; Rayner et al. 2019b; Pascoal et al. 2020), which raises the possibility that the *fw* genotype may be exposed to selection at an earlier stage than previously considered, for example through associated effects on foraging efficiency or predation risk. Further, it is possible that increased locomotion of *fw* juveniles might have accelerated the speed at which the mutation initially spread in the wild. Rather than facilitating local mating aggregations as we initially hypothesized, greater movement activity may instead permit silent crickets and females carrying male-silencing variants to encounter one another. The ultimate fitness consequences of these differences remain to be tested, but more broadly, this result illustrates how genetic correlations manifesting during development might impact the trajectory of a mutant genotype which carries fitness benefits at adult stages. The idea that advantageous mutations can have pleiotropic effects during development has been explored extensively in the context of insecticide resistance and alternative reproductive morphs (Boivin et al. 2001; Giraldo-Deck et al. 2020). Against the background of this work, our findings suggest the combined phenotypic and fitness effects of adaptive mutations arising from positive pleiotropy, negative pleiotropy, and genomic hitchhiking may be non-intuitive, and phenotypic variation caused by such effects can alter the dynamics of adaptive evolution as well as the manner in which plasticity affects that evolution.

In the crickets, transgenerational plasticity and within-generation plasticity sometimes acted in concert, sometimes in opposition, and in some cases did not appear influential in shaping trait variation. If TGP is adaptive for offspring (and therefore also for mothers), a match between parental and offspring environment should result in higher offspring fitness when compared to mis-matched offspring (Marshall and Uller 2007; Uller et al. 2013). However, we found no evidence supporting this expectation for offspring performance traits. Patterns of reproductive investment provide an illustrative example. Consistent with previous studies in this system and in other cricket species (Bailey et al. 2010; Conroy and Roff 2018), mothers who experienced No Song invested less in reproductive tissue. Such WGP is in line with adaptive predictions, as it allows individuals to reallocate resources to non-reproductive tissue when competition and/or opportunity for mating is low (Harshman and Zera 2007). This trade-off could be particularly adaptive in a flatwing-dominated social environment where no males can sing, because plasticity shifting resources from reproduction to survival could increase the chances that a female survives long enough to find a mate. Consistent effects of TGP and WGP on offspring would facilitate this, but instead we found that TGP and offspring WGP acted in opposing directions (Figure 3D). It is therefore unlikely that this TGP in reproductive investment is an adaptive, anticipatory effect to increase offspring fitness, but instead may be an incidentally-transmitted physiological consequence of mothers responding to their social environment (“selfish TGP” cf. Marshall and Uller 2007) or cross-generation spillover of parental condition (“condition-transfer effects” cf. Bonduriansky and Crean 2018). Lack of support for adaptive TGP is consistent with a recent meta-analysis which recovered weak evidence for it across taxa (Uller et al. 2013).

WGP has been suggested to be more efficient than TGP, so once capable of assessing their environment, offspring are expected to rewrite parental cues with their own (Ezard et al. 2014; Auge et al. 2017; Moore et al. 2019). Nevertheless, we found that the maternal social environment affected adult, but not juvenile, offspring phenotype in *T. oceanicus*. One reason for the delayed action of TGP might be that the maternal social environment influenced phenotype via mechanisms that would not cause observable differences until late in development. For example, many insects exhibit significant plasticity in the number of instars they undergo prior to sexual maturation, which can affect sexual size dimorphism at adulthood (Esperk et al. 2007; Stillwell et al. 2010). Another possibility is that the maternal social environment influenced offspring phenotype very early in development, such as size at hatching, but those effects dissipated prior to later phenotypic measurement, as was found in the salamander *Ambystoma talpoideum* (Moore et al. 2015). A third possibility is that TGP mediated by maternal social environments could be qualitatively different from TGP mediated by maternal physical environments. For example, nutritional or thermal environments that mothers experience may have more direct impacts on juvenile offspring, whereas the social environment comprised of adult social cues is likely to be of greater relevance to offspring when they are adults themselves.

In contrast to morphological traits, we found that behavioral traits related to movement (locomotive activity and likelihood of flight) were more influenced by WGP than TGP. This supports the prediction that traits whose expression remains flexible after development are more strongly affected by WGP (Beaty et al. 2016). Specifically, offspring of both sexes were more active when reared in song. They may increase walking activity to locate conspecifics they perceive to be abundant nearby, even in the absence of an immediate acoustic cue. In contrast, crickets raised without song have no indication of nearby conspecifics and may decrease short-range mate-searching via walking to instead wait for an acoustic cue. This trade-off is likely motivated by a high metabolic cost of mate-searching (Hack 1998) and the resulting increase in predation risk (Bell 1990). Our results contrast with previous studies in this species which found that adult males raised in song are less active than those raised in silence (Balenger and Zuk 2015) and that females exhibit limited flexibility in locomotive behavior in response to acoustic environment (Heinen-Kay et al. 2018), but are consistent with other findings that suggest increases in exploratory behavior under predation risk (Moschilla et al. 2021).

One explanation for this apparent inconsistency is that the latter studies conducted movement trials in environments that contained shelter or cover and in some cases assessed movement towards acoustic stimuli, whereas we used an open field test to mimic the experience of Hawaiian crickets within an all-flatwing population, in which locomotion in the absence of any immediately available acoustic signals is likely to have significant fitness consequences. Cover during movement trials could have reduced the perception of risk associated with walking, making increased undirected mate-searching in a song-less environment advantageous (Hedrick and Dill 1993). Additionally, two of the previous studies relied on indirect measures of activity (e.g., time spent walking, gridlines crossed, farthest grid reached), whereas we directly measured distance traveled using automated and validated coordinate collection, giving us greater resolution to resolve variation in activity. Finally, we made an incidental discovery during the course of the experiment which provides another intriguing explanation: offspring reared in No Song were more likely to attempt flight during the open field test, a pattern that was particularly strong in females who were ca. 4 times more likely to attempt fly if they had experienced no acoustic signals during development (Figure 4). Individuals reared without song may be more disposed to use flight as part of an un-directed, long-range dispersal strategy akin to Lévy flight (Viswanathan et al. 1996) to increase their chances of reaching an area of greater conspecific resources. Trading off increased long-range dispersal via flight with decreased walking behavior in song-less, flatwing-dominated environments could have increased the speed at which *flatwing* alleles spread under pressure from parasitoid flies.

**Figure 4.**
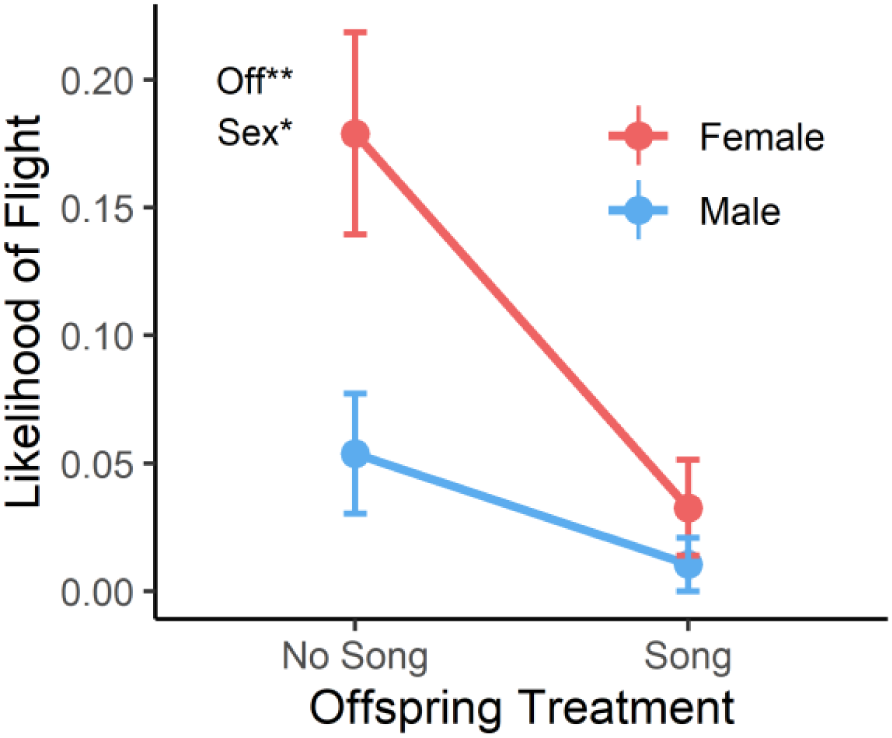
Likelihood of flight during open field trials. Means and ± 1 standard error are represented by circles and bars, respectively. Significance of offspring (“Off”) treatment and sex are indicated with asterisks: \**p* <0.05, \*\**p* <0.01.

Our results support the idea that the effects of TGP can be contingent upon offspring environment. Put another way, sometimes TGP potentiates WGP, and sometimes it suppresses WGP. It is therefore necessary to consider the potentially conflicting effects of WGP and TGP when predicting how phenotypic plasticity influences adaptive evolution. Theory predicts that, following rapid environmental change, populations may exhibit a transient increase in plasticity because genotypes which shift trait expression closer to a new optimum are favored (Lande 2009). Additionally, genotypes coding for reaction norm slopes of other traits that offset negative effects of new variants spreading under selection may also be favored, as appears to be the case in *T. oceanicus* in Hawaii and other systems (Bailey et al. 2021). If increases in WGP during an adaptive evolutionary response result in a spillover of non-adaptive TGP to the offspring generation, the role of plasticity in facilitating the establishment and spread of novel adaptations may be less straightforward than currently understood (Lacey 1998; Bonduriansky and Day 2009; Bell and Hellmann 2019). Previous work in the Hawaiian flatwing cricket system supports predictions of increased WGP which mitigates negative pleiotropy of male silence, in this case facilitating mate location and reproduction in a song-less social environment (Bailey and Zuk 2008; Bailey et al. 2018; Tinghitella et al. 2009; Bailey 2011; Bailey and Zuk 2012; Balenger and Zuk 2015; Pascoal et al. 2018), but see (Rayner et al. 2020). However, we did not find a prominent signature of either adaptive TGP or of an adaptive interaction between TGP and WGP. Instead, our results suggest that plasticity transmitted across generations may have more complex effects and, in some cases, counterbalance facilitating effects of WGP.

## Acknowledgements

We are grateful for support from the Natural Environment Research Council to NWB (NE/L011255/1 and NE/T000619/1). T. Hitchcock patiently tolerated the execution of a cricket experiment in a converted spare room of his flat during the first 2020 UK Covid-19 lockdown. We are grateful for the assistance of D. Forbes, A. Grant and M. McGunnigle in cricket rearing and laboratory maintenance. S. Pascoal advised on technical aspects of the social environment manipulation. J.G. Rayner provided valuable feedback that improved the experimental design, and T.M. Jones and L.E. Rendell gave helpful input on the writing and interpretation of results.

