## Supplemental_Info for "Within-generation and transgenerational social plasticity interact during rapid adaptive evolution"

#### Contents

Supplementary Methods………………………………………………………………………… S2

Supplementary Results……………………………………………………………………… S6

Supplementary Tables………………………………………………………………………… S7

Supplementary Figures……………………………………………………………………… S18

### Supplementary Methods

#### DORIS coordinate collection path smoothing and external validation

Initial trials using DORIS tracking software revealed a bug in which the software often recorded a series of very small movements across the length of the cricket rather than one static point, even when crickets were motionless. This issue was fixed in R by slightly smoothing the coordinate paths. We then externally validated smoothing methods using a subset of video clips. For coordinate smoothing, we averaged x and y coordinates collected from DORIS software over 24 frames to create a smoothed path, then removed coordinate pairs which were less than 1 mm from the position in the previous frame. To externally validate this fix and ensure that our distance calculation method was accurate, we randomly selected 100-second increments from 8 videos, extracted one JPEG for every 10 frames of the video, then manually measured the path across these images in ImageJ (v.1.8.0_112). These manually collected coordinates produced distance values that were very highly correlated with distance values based on smoothed-DORIS coordinates (Pearson correlation: $r^2 = 0.999$, $n = 8$, $p < 0.001$), providing confidence in the validity of the smoothing methods (Figure S1).

#### Measurement and Statistical Analysis of Other Movement Parameters

We calculated proportion of the arena entered as a measure of exploratory activity ("*proportion explored*") by overlaying coordinate data onto a 11x17 cm raster grid containing 187 distinct 1 $cm^2$ squares, then summing how many squares the cricket entered and dividing by the total number of non-origin squares (186). We calculated time spent in three zones: "*origin time*" (the square where the individual started the test), "*edge time*" (10 mm border around the perimeter of the arena), and "*middle time*" (all other space in the arena). This enabled us to investigate differences in space usage, under the assumption that greater edge usage (thigmotaxis) represents less risky movement behavior. However, for Experiment 3 which involved adult crickets in a larger arena, when calculating *edge time* and *middle time* and *origin time*, we divided the arena into a raster of 1,517 distinct 1 $cm^2$ squares. Due to the larger size of adult crickets and larger arena, the 'edge' region was defined as a 2 $cm^2$ border along the perimeter, the 'origin' was defined as the 1 $cm^2$ grid in which the cricket began its trial, and the 'middle' region was all other space not defined as edge or origin.

In Experiment 1, we compared *proportion explored, origin time, middle time,* and *edge time* between wing morph genotypes in 15-day old and 45-day old offspring using separate linear models. *Distance* and *proportion explored* were strongly correlated (Pearson correlation: $r^2 = 0.709$, $n = 254$ $p < 0.001$ for 15-day old nymphs and $r^2 = 0.846$, $n = 224$ $p < 0.001$ for 45-day nymphs). Models examining *proportion explored* were run both with and without *distance* as a covariate to check whether any genotypic effects on *proportion explored* were explained by variation in distance. With the exception of the analysis of *origin time*, sex did not approach significance in any of these models (all $p > 0.2$) and the qualitative outcomes did not differ. Thus, models retaining all individuals, and excluding sex as a fixed effect, were retained except in the case of *origin time* (Table S2).

In Experiment 3, we examined the effect of maternal treatment on juvenile offspring locomotive activity using separate LMMs for: *proportion explored, edge time*, *middle time,* and *origin time*. All models included maternal treatment and experimental replicate as categorical factors; temperature, time of day, and pronotum length as covariates; and maternal ID as a random effect. The treatment*replicate interaction ($p > 0.2$) was excluded from the final models. Similar models were run for 45-day old offspring, except experimental replicate was not included because we only had 45-day data for trial 1. The general form of the models is given by Equation 6 in Table S1. For adult offspring in Experiment 3, we then tested the impact of TGP and WGP on adult locomotive behaviors using separate LMMs for each behavior and sex. All models included maternal treatment, offspring treatment, temperature, time, and somatic SMI, plus box replicate as a random effect (Equation 8 in Table S1). Non-significant ($p > 0.2$) maternal*offspring treatment interactions were removed. As in Experiment 1, we modelled proportion of grid explored both with and without distance as a covariate.

#### Acoustic treatments using in Experiment 2 and 3

To create male song playback used for the Song treatment, .WAV files were made by recording 24 Kauai males in a controlled laboratory setting, calculating average song parameters, and constructing two artificial calling files with these averaged parameters (Pascoal et al. 2018). Here, the two calls were each played for 1 hr sequentially at the beginning and end of the night cycle, when calling naturally peaks in the wild (8:00-10:00

and 18:00-20:00). In the middle of the night cycle (10:00-18:00), we played one call at a time at 15-minute intervals, with five minutes of silence between each alternating call.

#### Experiment 3 procedures during 2020 Covid-19 lockdown

To limit necessary maintenance for technicians with "essential worker" status, we pooled egg pads from Experiment 3 into five replicate 16-L boxes per maternal acoustic treatment. Each box contained 6-7 egg pads (each egg pad from one pair of adults). We organized the pooling such that egg pads in Silent Box 1 were collected from matings that occurred at the same dates as those that produced egg pads in Song Box 1, so that offspring from Silent Box 1 and Song Box 1 hatched at the same time. Thus, crickets in Silent Box 1 and Song Box 1 were older than crickets in Silent Box 2 and Song Box 2 and so on. These 10 boxes were initially housed in the general incubator at 25°C for two weeks after the lockdown closure. After two weeks, the university granted us permission to bring the crickets back to a private apartment to relieve the workload of technicians, where we kept them in an undisturbed spare room. The same level of precision in temperature-regulation as in the lab incubators could not be achieved. The room was kept at approximately 18-24°C throughout their development. The experimental blocks we ran across the pandemic disruption were always paired, such that subjects experiencing Song and No Song treatments were exposed to the same potential block effects over time. To control for any statistical variation arising from this, we included a box replicate term in all models. For example, individuals from Song Box 1 and No Song Box 1 belonged to box replicate 1, and so on. Because boxes of the same box replicate hatched at the same time and therefore experienced any potential changes in rearing temperature at the same approximate stage in development, including this term in models controlled for any inconsistencies in environmental factors during rearing. The window in the room was blocked out using emergency blankets, and two lamps were plugged into timer-controlled outlets to match the photo-reversed light:dark cycle set in the general incubator.

Egg pads still containing un-hatched eggs at this time were placed in separate containers (to keep track of sibling relationships). We isolated hatchlings and kept them at 18-24 °C on a 12h:12h photo-reversed light:dark cycle for their entire development. Before we could bring these crickets to the apartment, most eggs had already hatched, limiting the number of individuals available for the early juvenile offspring TGP experiment. Therefore, to supplement this original data, the experiment was repeated approximately two months later once the lab had re-opened. All rearing conditions were the same in this second trial, except individuals were kept at a constant 25° in an incubator in the laboratory. Individuals which had already hatched before permission was granted to take them back to the apartment were kept in their original pooled boxes in apartment rearing conditions and later used for the adult offspring TGP experiment. Once we were allowed back into the laboratory, we returned the 10 boxes to the general incubator. These slight rearing temperature fluctuations were unavoidable. However, because crickets were switched to warmer conditions at different stages in development, and box replicates of crickets were spread across both treatments evenly (see Main Text), we minimized the risk that these fluctuations systematically biased the experiment. We additionally modelled effects of Trial 1 vs. Trial 2 to account for environmental differences during lockdown vs. during normal operation of the lab, and we also included age class as a random factor in all models (see Experiment 3, *Statistical Analyzes* in the Main Text). For juvenile offspring, OFTs were performed at 15 and 45 days in Trial 1 of this experiment in the spare room of the private apartment at 23-25.5°C. For Trial 2 of the experiment, the OFTs were conducted in the same incubator as in Experiment 1 at 24-28°C constraints, during trial 2. Temperature variation during the OFT was accounted for in statistical models described in the Main Text.

### Supplementary Results

#### Experiment 1: Genotypic differences in juvenile locomotion

Homozygous *fw* individuals at both ages showed different patterns of exploratory behavior: they entered a greater proportion of the arena, spent more time along the edge of the arena, and spent less time at the origin than homozygous *nw* individuals (Table S8, Figure S3, Figure S6). However, in all models of proportion of grids explored, when distance was included as a covariate there were no longer a significant effect of morph (Table S9). This indicates that although *fw*-carrying individuals explore more grid squares, this difference may be associated with their greater movement distances as opposed to more exploratory behavior *per se*.

#### Experiment 3: TGP, WGP, and their interaction

TGP affected the proportion of time adult female offspring spent in the middle of the test arena – a proxy for exploratory behavior (Supplemental Table S10, Figure S7).

### Supplementary Tables

**Table S1.** Transformations applied to response variables to satisfy assumptions of normality

| | ***Variable*** | ***Transformation applied*** |
|---|---|---|
| Experiment 1 | *Distance at* 15 days | square-root |
| | *Proportion of grids explored* at 15 days | square-root |
| | *Edge time* at 15 days | none |
| | *Middle time* at 15 days | natural log |
| | *Origin time* at 15 days | natural log |
| | *Distance at* 45 days | natural log |
| | *Proportion of grids explored* at 45 days | square-root |
| | *Edge time* at 45 days | square-root |
| | *Middle time* at 45 days | none |
| | *Origin time* at 45 days | square-root |
| Experiment 2 | *SMI* | natural log |
| | *Reproductive tissue weight* | natural log |
| | *Likelihood of mounting* | none |
| | *Likelihood of spermatophore transfer* | none |
| Experiment 3 | *Size* at 15 days | square-root |
| | *Distance* at 15 days | none |
| | *Proportion of grids explored* at 15 days | none |
| | *Edge time* at 15 days | none |
| | *Middle time* at 15 days | square-root |
| | *Origin time* at 15 days | natural log |
| | *Size at 45 days* | square-root |
| | *Distance at 45 days* | none |
| | *Proportion of grids explored* at 45 days | square-root |
| | *Edge time* at 45 days | square-root |
| | *Middle time* at 45 days | square-root |
| | *Origin time* at 45 days | square-root |
| | *Size* in adult females | natural log |
| | *Reproductive tissue mass* in adult females | natural log |
| | *Somatic SMI* in adult females | square-root |
| | *Distance* in adult females | square-root |
| | *Proportion of grids explored* in adult females | natural log |
| | *Edge time* in adult females | square-root |
| | *Middle time* in adult females | square-root |
| | *Origin time* in adult females | square-root |
| | *Size* in adult males | none |
| | *Reproductive tissue mass* in adult males | none |
| | *Somatic SMI* in adult males | natural log |
| | *Distance* in adult males | square-root |
| | *Proportion of grids explored* in adult males | none |
| | *Edge time* in adult males | square-root |
| | *Middle time* in adult males | square-root-transformed |
| | *Origin time* in adult males | square-root-transformed |

**Table S2.** Equations describing general forms of statistical models for each experiment

| | |
|---|---|
| Experiment 1 | **Eqn. 1** *movement response ~ intercept + morph + sex + morph:line + pronotum length + temperature + time of day +* $\varepsilon$ |
| Experiment 2 | **Eqn. 2** *female SMI ~ intercept + acoustic treatment + days isolated + replicate +* $\varepsilon$ <br> **Eqn. 3** *mating behavior ~ intercept + acoustic treatment + female SMI + male SMI +* $\varepsilon$ <br> **Eqn. 4** *ovary mass ~ intercept + acoustic treatment + days isolated + soma mass +* $\varepsilon$ |
| Experiment 3 | **Eqn. 5** *juvenile offspring size ~ intercept + maternal treatment + replicate + maternal treatment * replicate + (1* $\mid$ *maternal ID) +* $\varepsilon$ <br> **Eqn. 6** *movement response ~ intercept + maternal treatment + replicate + temperature + time +* $\varepsilon$ <br> **Eqn. 7** *adult offspring trait ~ intercept + maternal treatment + offspring treatment + maternal treatment * offspring treatment + (1* $\mid$ *age class) + pronotum length + soma mass +* $\varepsilon$ <br> **Eqn. 8** *movement response ~ intercept + maternal treatment + offspring treatment + maternal * offspring treatment + temperature + time + soma mass + (1* $\mid$ *age class) +* $\varepsilon$ <br> **Eqn. 9** *flight attempts ~ intercept + maternal treatment + offspring treatment + sex + maternal treatment * sex + somatic SMI + (1* $\mid$ *age class) +* $\varepsilon$ |

Optional scaling covariates shown in grey text

**Table S3.** Linear models examining area of arena explored for 15 day-old (top) and 45 day-old (bottom) juveniles, excluding individuals that remained stationary throughout the test

| | | Proportion explored | | | Edge time | | | Middle time | | | Origin time | | |
|---|---|---|---|---|---|---|---|---|---|---|---|---|---|
| | | ***df*** [*] | ***F*** | ***P*** [†] | ***df*** | ***F*** | ***P*** | ***df*** | ***F*** | ***P*** | ***df*** | ***F*** | ***P*** |
| 15 days old | Morph | 1, 222 | 12.244 | **<0.001** | 1, 222 | 18.879 | **<0.001** | 1, 222 | 0.405 | 0.525 | 1, 188 | 20.465 | **<0.001** |
| | Morph:Line | 4, 222 | 3.613 | 0.007 | 4, 222 | 0.576 | 0.681 | 4, 222 | 2.127 | 0.078 | 4, 188 | 1.561 | 0.186 |
| | Pronotum | 1, 222 | 1.261 | 0.363 | 1, 222 | 0.721 | 0.397 | 1, 222 | 3.660 | 0.057 | 1, 188 | 2.622 | 0.107 |
| | Temperature | 1, 222 | 2.422 | 0.121 | 1, 222 | 0.910 | 0.341 | 1, 222 | 2.628 | 0.106 | 1, 188 | 0.136 | 0.713 |
| | Assay time | 1, 222 | 2.310 | 0.130 | 1, 222 | 0.011 | 0.915 | 1, 222 | 0.663 | 0.416 | 1, 188 | 0.246 | 0.620 |
| | Sex | - | - | - | - | - | - | - | - | - | 1, 188 | 3.191 | 0.076 |
| 45 days old | Morph | 1, 156 | 8.383 | **0.004** | 1, 156 | 6.277 | **0.013** | 1, 156 | 0.230 | 0.632 | 1, 156 | <0.001 | 0.989 |
| | Morph:Line | 4, 156 | 2.361 | 0.056 | 4, 156 | 5.258 | **<0.001** | 4, 156 | 0.698 | 0.594 | 4, 156 | 3.043 | 0.019 |
| | Pronotum | 1, 156 | 7.889 | **0.006** | 1, 156 | 15.634 | **<0.001** | 1, 156 | 0.001 | 0.975 | 1, 156 | 7.108 | 0.008 |
| | Temperature | 1, 156 | 0.029 | 0.865 | 1, 156 | 1.198 | 0.276 | 1, 156 | 0.073 | 0.788 | 1, 156 | 1.908 | 0.169 |
| | Assay time | 1, 156 | 29.679 | **<0.001** | 1, 156 | 2.609 | 0.108 | 1, 156 | 0.961 | 0.328 | 1, 156 | 0.189 | 0.664 |

[*] numerator, denominator
[†] significance at α<0.05 indicated in bold

**Table S4.** Counts of individuals used in adult TGP offspring experiment (Experiment 3) by sex and maternal – offspring acoustic treatment pairings.

| Treatment | Female | Male | Total |
|---|---|---|---|
| Maternal NoSo* – Offspring NoSo | 48 (46)† | 47 | 95 (93) |
| Maternal NoSo – Offspring So | 46 | 50 | 96 |
| Maternal So – Offspring NoSo | 49 (48) | 46 | 95 (94) |
| Maternal So – Offspring So | 47 (46) | 46 (45) | 93 (91) |
| | 190 (186) | 189 (188) | 379 (374) |

*NoSo = No Song acoustic treatment; So = Song treatment.
†Numbers in parentheses indicate the count of individuals with open field test data, if different from the count with morphological data. I.e., 48 (46) describes 48 individuals with morphological data but only 46 with both morphological data and open field test data.

**Table S5.** Generalized linear models examining the effects of WGP arising from the acoustic rearing environment on female mating behavior in the maternal generation

| | **Mounting** | | | **Spermatophore Transfer** | | |
|---|---|---|---|---|---|---|
| | ***df*** * | $\chi^2$ | ***P*** † | ***df*** | $\chi^2$ | ***P*** |
| Acoustic treatment | 1,61 | 0.340 | 0.560 | 1,45 | 0.332 | 0.565 |
| Male condition | 1,61 | 6.789 | **0.009** | 1,45 | 2.864 | 0.091 |
| Female condition | 1,61 | 2.772 | 0.096 | 1,45 | 5.361 | **0.021** |

* numerator, denominator

† $p < 0.05$ indicated in bold

**Table S6.** Linear models examining effects of TGP arising from the maternal acoustic environment on proportions of the test arena explored and time spent in each area of the arena by 15 day-old (top) and 45 day-old (bottom) juveniles

| | | Proportion explored | | | Edge time | | | Middle time | | | Origin time | | |
|---|---|---|---|---|---|---|---|---|---|---|---|---|---|
| | | ***df***[*] | ***F*** | ***P***[†] | ***df*** | ***F*** | ***P*** | ***df*** | ***F*** | ***P*** | ***df*** | ***F*** | ***P*** |
| 15 days old | Maternal treatment | 1, 299 | 0.156 | 0.693 | 1, 300 | 0.267 | 0.605 | 1, 300 | 0.801 | 0.370 | 1, 300 | 0.514 | 0.473 |
| | Experimental replicate | 1, 299 | 22.208 | **<0.001** | 1, 300 | 0.127 | 0.721 | 1, 300 | 3.959 | **0.047** | 1, 300 | 0.012 | 0.913 |
| | Pronotum | 1, 299 | 0.034 | 0.854 | 1, 300 | 5.699 | **0.017** | 1, 300 | 2.492 | 0.114 | 1, 300 | 1.909 | 0.167 |
| | Temperature | 1, 299 | 0.062 | 0.804 | 1, 300 | 1.856 | 0.173 | 1, 300 | 0.359 | 0.549 | 1, 300 | 2.235 | 0.135 |
| | Assay time | 1, 299 | 3.136 | 0.077 | 1, 300 | 0.245 | 0.620 | 1, 300 | 2.245 | 0.134 | 1, 300 | 4.102 | **0.043** |
| | Mat. treatment* Exp. replicate | 1, 299 | 2.892 | 0.089 | - | - | - | - | - | - | - | - | - |
| 45 days old | Maternal treatment | 1, 195 | 0.541 | 0.462 | 1, 195 | 0.147 | 0.701 | 1, 195 | <0.001 | 0.992 | 1, 195 | 0.323 | 0.570 |
| | Pronotum | 1, 195 | 2.987 | 0.085 | 1, 195 | 0.454 | 0.500 | 1, 195 | 0.255 | 0.614 | 1, 195 | 0.046 | 0.831 |
| | Temperature | 1, 195 | 5.695 | **0.017** | 1, 195 | 2.422 | 0.120 | 1, 195 | 0.839 | 0.360 | 1, 195 | 3.895 | 0.048 |
| | Assay time | 1, 195 | 2.764 | 0.096 | 1, 195 | 0.128 | 0.720 | 1, 195 | 4.402 | **0.036** | 1, 195 | 0.163 | 0.687 |

[*] numerator, denominator
[†] significance at α<0.05 indicated in bold

**Table S7.** Linear models examining effects of TGP arising from the maternal acoustic environment on pronotum length of 15 day-old (top) and 45 day-old (bottom) juveniles

| | | ***df**** | $\chi^2$ | ***P***† |
|---|---|---|---|---|
| 15 days old | Maternal treatment | 1, 307 | 0.046 | 0.829 |
| | Experimental replicate | 1, 307 | 232.953 | **<0.001** |
| 45 days old | Maternal treatment | 1, 198 | 0.732 | 0.392 |

* numerator, denominator

† significance at α<0.05 indicated in bold

**Table S8.** Linear models examining areas of arena explored by 15 day-old (top) and 45 day-old (bottom) juveniles in open field tests

| | | Proportion explored | | | Edge time | | | Middle time | | | Origin time | | |
|---|---|---|---|---|---|---|---|---|---|---|---|---|---|
| | | ***df***[*] | ***F*** | ***P***[†] | ***df*** | ***F*** | ***P*** | ***df*** | ***F*** | ***P*** | ***df*** | ***F*** | ***P*** |
| 15 days old | Morph | 1, 245 | 11.110 | **<0.001** | 1, 245 | 18.213 | **<0.001** | 1, 245 | 0.902 | 0.343 | 1, 206 | 17.731 | **<0.001** |
| | Morph:Line | 4, 245 | 3.784 | **0.005** | 4, 245 | 1.162 | 0.328 | 4, 245 | 2.680 | **0.032** | 4, 206 | 2.385 | 0.052 |
| | Pronotum | 1, 245 | 0.134 | 0.714 | 1, 245 | 1.127 | 0.261 | 1, 245 | 0.663 | 0.416 | 1, 206 | 3.130 | 0.078 |
| | Temperature | 1, 245 | 2.267 | 0.133 | 1, 245 | 0.554 | 0.457 | 1, 245 | 2.161 | 0.143 | 1, 206 | 0.166 | 0.685 |
| | Assay time | 1, 245 | 1.676 | 0.197 | 1, 245 | 0.020 | 0.887 | 1, 245 | 0.423 | 0.032 | 1, 206 | 0.181 | 0.671 |
| | Sex | - | - | - | - | - | - | - | - | - | 1, 206 | 4.107 | **0.044** |
| 45 days old | Morph | 1, 210 | 14.336 | **<0.001** | 1, 210 | 12.196 | **<0.001** | 1, 210 | 0.498 | 0.481 | 1, 210 | 1.426 | 0.234 |
| | Morph:Line | 4, 210 | 8.920 | **<0.001** | 4, 210 | 12.577 | **<0.001** | 4, 210 | 4.437 | **0.002** | 4, 210 | 10.533 | **<0.001** |
| | Length | 1, 210 | 10.833 | **0.001** | 1, 210 | 17.707 | **<0.001** | 1, 210 | 1.285 | 0.258 | 1, 210 | 10.285 | **0.002** |
| | Temperature | 1, 210 | 0.264 | 0.608 | 1, 210 | 1.809 | 0.180 | 1, 210 | 0.108 | 0.743 | 1, 210 | 2.222 | 0.138 |
| | Assay Time | 1, 210 | 44.405 | **<0.001** | 1, 210 | 12.454 | **<0.001** | 1, 210 | 5.667 | **0.018** | 1, 210 | 4.492 | **0.035** |

[*] numerator, denominator
[†] significance at α<0.05 indicated in bold

**Table S9.** Linear models examining area of arena explored by 15 day-old (top) and 45 day-old (bottom) juveniles in open field tests, with *distance* included as a covariate

| | | Proportion explored | | | Edge time | | | Middle time | | | Origin time | | |
|---|---|---|---|---|---|---|---|---|---|---|---|---|---|
| | | ***df***[*] | ***F*** | ***P***[†] | ***df*** | ***F*** | ***P*** | ***df*** | ***F*** | ***P*** | ***df*** | ***F*** | ***P*** |
| 15 days old | Morph | 1, 244 | 0.002 | 0.968 | 1, 244 | 5.969 | **0.015** | 1, 244 | 0.386 | 0.535 | 1, 205 | 12.374 | **<0.001** |
| | Morph:Line | 4, 244 | 1.846 | 0.121 | 4, 244 | 0.359 | 0.837 | 4, 244 | 1.379 | 0.242 | 4, 205 | 0.628 | 0.643 |
| | Pronotum | 1, 244 | 0.188 | 0.665 | 1, 244 | 4.241 | **0.041** | 1, 244 | 0.389 | 0.534 | 1, 205 | 9.337 | **0.003** |
| | Temperature | 1, 244 | 3.309 | 0.070 | 1, 244 | 3.055 | 0.082 | 1, 244 | 1.533 | 0.217 | 1, 205 | 0.947 | 0.332 |
| | Assay time | 1, 244 | 3.601 | 0.059 | 1, 244 | 0.098 | 0.754 | 1, 244 | 0.215 | 0.644 | 1, 205 | 0.993 | 0.320 |
| | Distance | 1, 244 | 11182.289 | **<0.001** | 1, 244 | 201.053 | **<0.001** | 1, 244 | 47.965 | **<0.001** | 1, 205 | 170.915 | **<0.001** |
| | Sex | - | - | - | - | - | - | - | - | - | 1, 205 | 3.778 | **0.053** |
| 45 days old | Morph | 1, 209 | 0.061 | 0.806 | 1, 209 | 2.586 | 0.109 | 1, 209 | 3.466 | 0.064 | 1, 209 | 7.536 | **0.007** |
| | Morph:Line | 4, 209 | 1.475 | 0.211 | 4, 209 | 6.667 | **<0.001** | 4, 209 | 0.495 | 0.740 | 4, 209 | 1.124 | 0.346 |
| | Length | 1, 209 | 0.033 | 0.857 | 1, 209 | 7.369 | 0.007 | 1, 209 | 0.945 | 0.332 | 1, 209 | 0.862 | 0.354 |
| | Temperature | 1, 209 | 0.220 | 0.639 | 1, 209 | 1.982 | **0.161** | 1, 209 | 0.013 | 0.910 | 1, 209 | 1.907 | 0.169 |
| | Assay time | 1, 209 | 10.774 | **0.001** | 1, 209 | 1.689 | 0.195 | 1, 209 | 0.928 | 0.336 | 1, 209 | 8.262 | **0.004** |
| | Distance | 1, 209 | 1235.053 | **<0.001** | 1, 209 | 216.583 | **<0.001** | 1, 209 | 97.169 | **<0.001** | 1, 209 | 244.856 | **<0.001** |

[*] numerator, denominator
[†] significance at α<0.05 indicated in bold

**Table S10.** Linear models examining the effects of TGP arising from the maternal acoustic environment and WGP arising from the offspring acoustic environment on proportions of the test area explored by adult females (top) and adult males (bottom)

| | | Proportion explored | | | Edge time | | | Middle time | | | Origin time | | |
|---|---|---|---|---|---|---|---|---|---|---|---|---|---|
| | | ***df*** [*] | ***F*** | ***P*** [†] | ***df*** | ***F*** | ***P*** | ***df*** | ***F*** | ***P*** | ***df*** | ***F*** | ***P*** |
| 15 days old | Maternal treatment | 1, 181 | 3.563 | 0.059 | 1, 181 | 0.203 | 0.652 | 1, 180 | 5.580 | **0.018** | 1, 181 | 0.031 | 0.861 |
| | Offspring treatment | 1, 181 | 6.080 | **0.014** | 1, 181 | 2.645 | 0.104 | 1, 180 | 1.380 | 0.240 | 1, 181 | 2.808 | 0.094 |
| | Temperature | 1, 181 | 1.637 | 0.201 | 1, 181 | 1.612 | 0.204 | 1, 180 | 1.513 | 0.219 | 1, 181 | 6.978 | **0.008** |
| | Assay time | 1, 181 | 1.487 | 0.223 | 1, 181 | 2.019 | 0.155 | 1, 180 | 1.152 | 0.283 | 1, 181 | 5.236 | **0.022** |
| | Somatic condition | 1, 181 | 0.856 | 0.355 | 1, 181 | 4.360 | **0.037** | 1, 180 | 0.437 | 0.508 | 1, 181 | 2.332 | 0.127 |
| | Maternal treatment * offspring treatment | - | - | - | - | - | - | 1, 180 | 5.661 | **0.017** | - | - | - |
| 45 days old | Maternal treatment | 1, 183 | 0.291 | 0.590 | 1, 183 | 0.211 | 0.646 | 1, 183 | 0.114 | 0.736 | 1, 183 | <0.001 | 0.998 |
| | Offspring treatment | 1, 183 | 3.258 | 0.071 | 1, 183 | 2.056 | 0.152 | 1, 183 | 2.297 | 0.130 | 1, 183 | 0.108 | 0.743 |
| | Temperature | 1, 183 | 5.383 | **0.020** | 1, 183 | 1.533 | 0.216 | 1, 183 | 0.006 | 0.940 | 1, 183 | 0.630 | 0.428 |
| | Assay time | 1, 183 | 2.059 | 0.151 | 1, 183 | 0.244 | 0.621 | 1, 183 | 0.059 | 0.808 | 1, 183 | 1.980 | 0.159 |
| | Somatic condition | 1, 183 | 3.508 | 0.061 | 1, 183 | 0.005 | 0.942 | 1, 183 | 0.284 | 0.594 | 1, 183 | 0.414 | 0.520 |

[*] numerator, denominator

[†] significance at α<0.05 indicated in bold

### Supplementary Figures

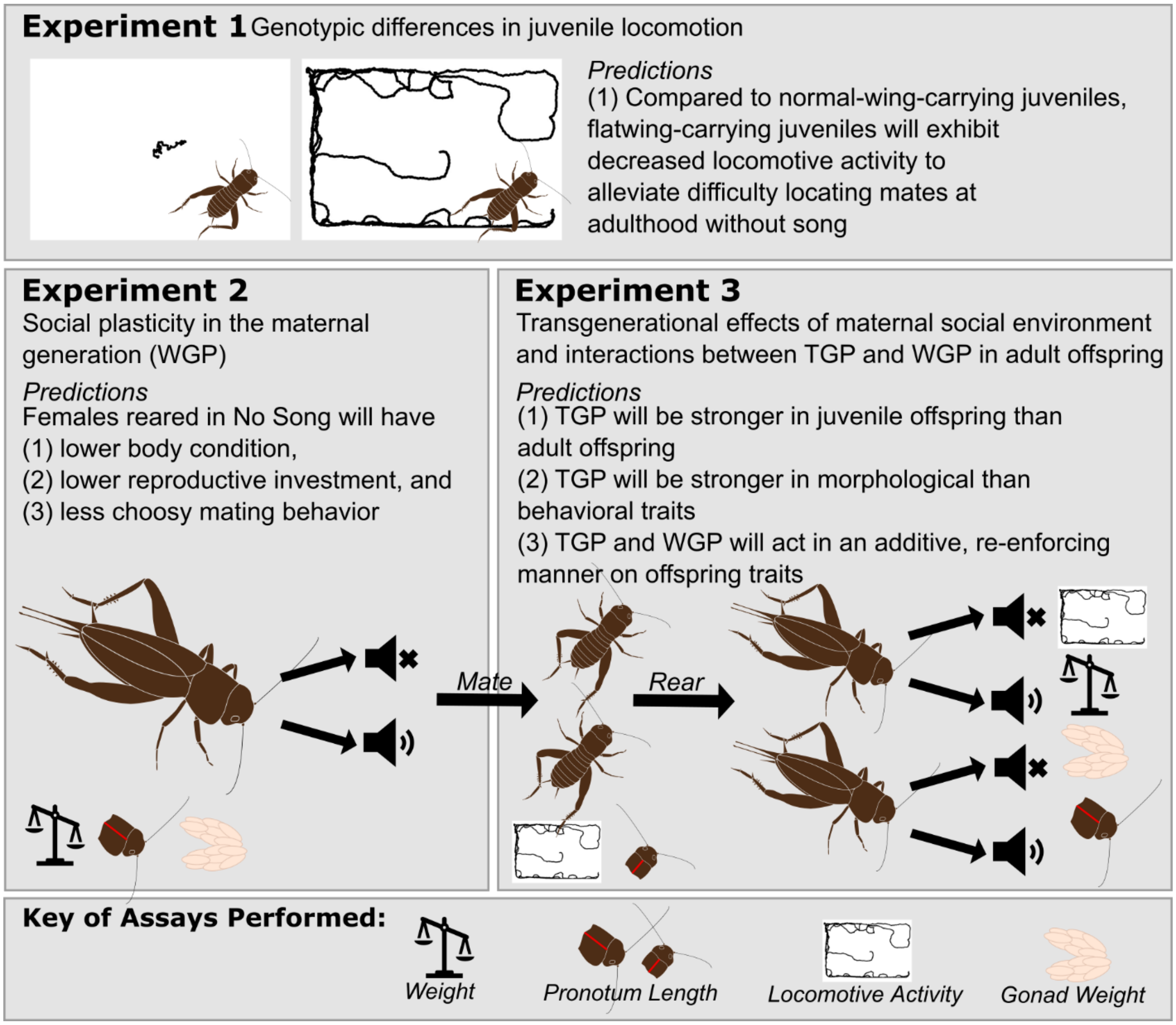

**Figure S1.** Description of three experiments performed to test whether and how interactions among genotypic, within-generation plasticity (WGP), and transgenerational plasticity (TGP) facilitate rapid adaptive evolution, using silent Hawaiian field crickets *Teleogryllus oceanicus* as a model system. Predicted effects of genotypic, WGP and TGP are given for each experiment, and diagrams depict assays performed to test effects on key fitness traits known to influence the spread of silent crickets: condition, locomotion, and reproductive tissues.

151

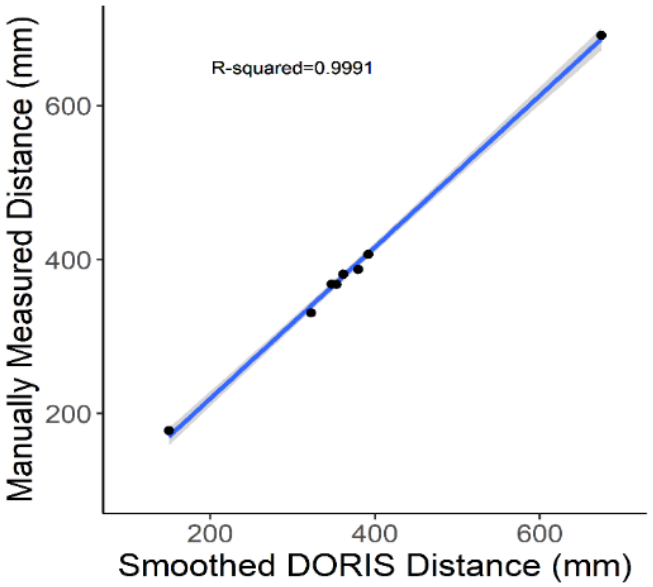

**Figure S2.** Validation test examining relationship between smoothed DORIS-calculated distance values and manually measured distance values in 8 videos.

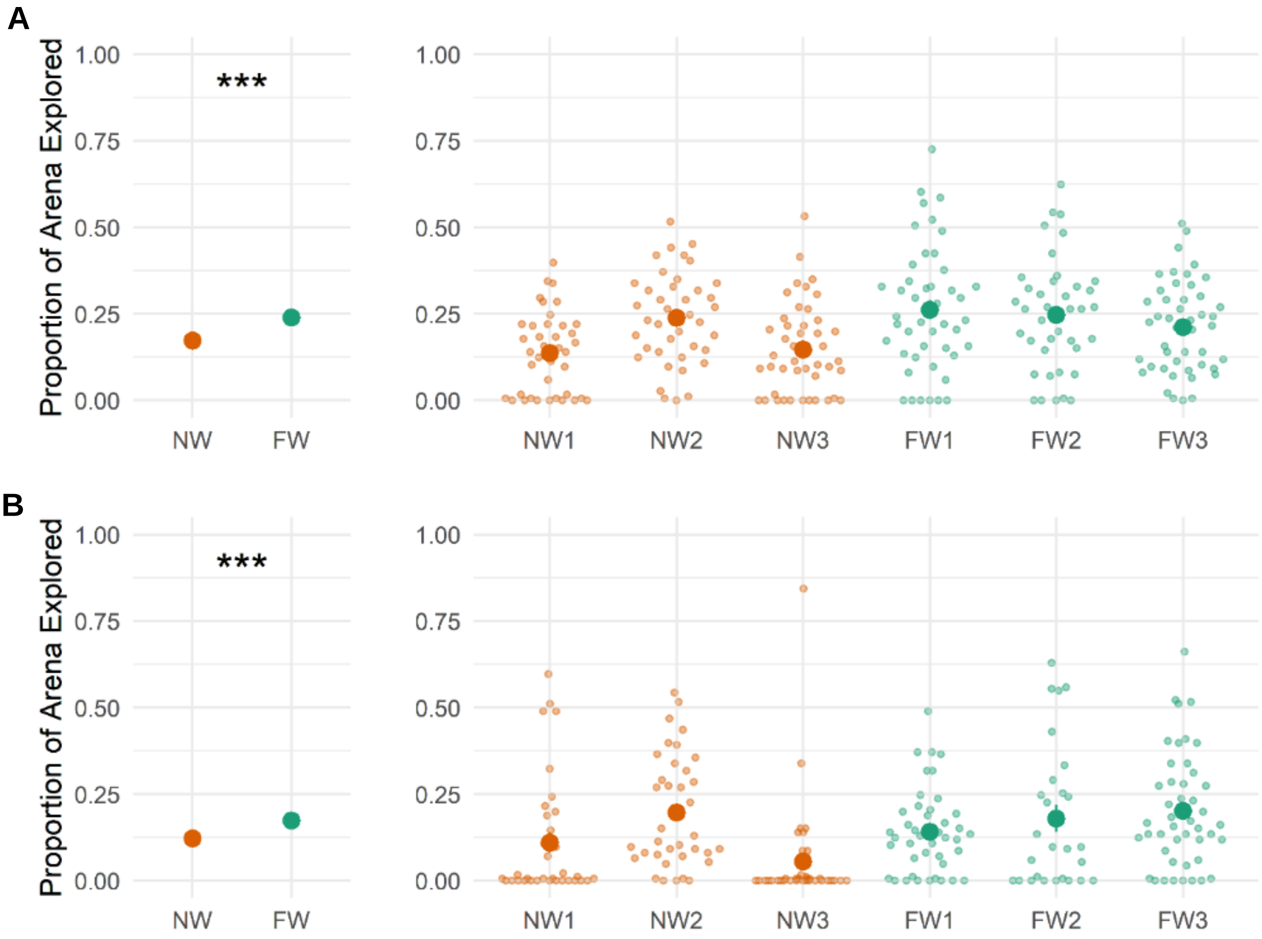

**Figure S3.** The effect of wing morph genotype on proportion of the arena explored by juveniles in open field tests for (**A**) 45 day-old nymphs and (**B**) 45 day-old nymphs. Plots on the left illustrate pooled means across three replicate morph lines. Bars indicating ± 1 standard error are not shown as these are too small to indicate graphically without being obstructed by the symbols for means. Violin plots on the right show data for each replicate morph line, with dark circles indicating means, small light circles showing each data point, and bars indicating ± 1 standard error (when visible). *** indicates a morph difference with a p-value < 0.001.

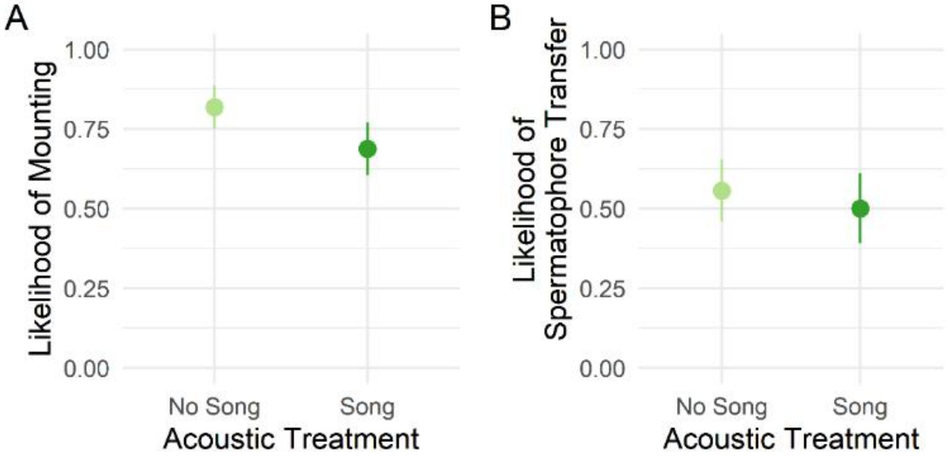

**Figure S4.** The effect of acoustic environment on (**A**) likelihood of mounting, and (**B**) likelihood of receiving a spermatophore given the occurrence of mounting. All plots show means ± 1 standard error.

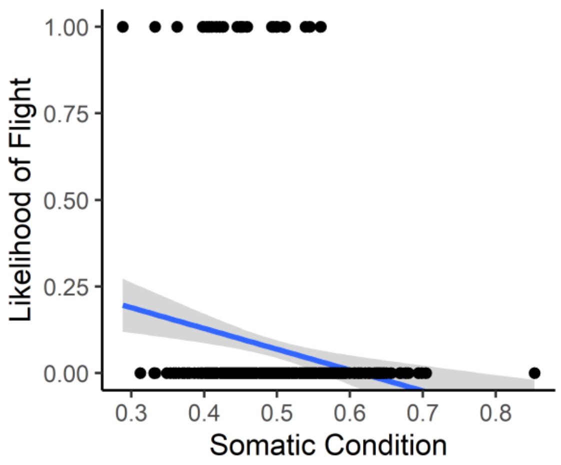

**Figure S5**. The relationship between somatic condition and flight probability in adult offspring during the open field test.

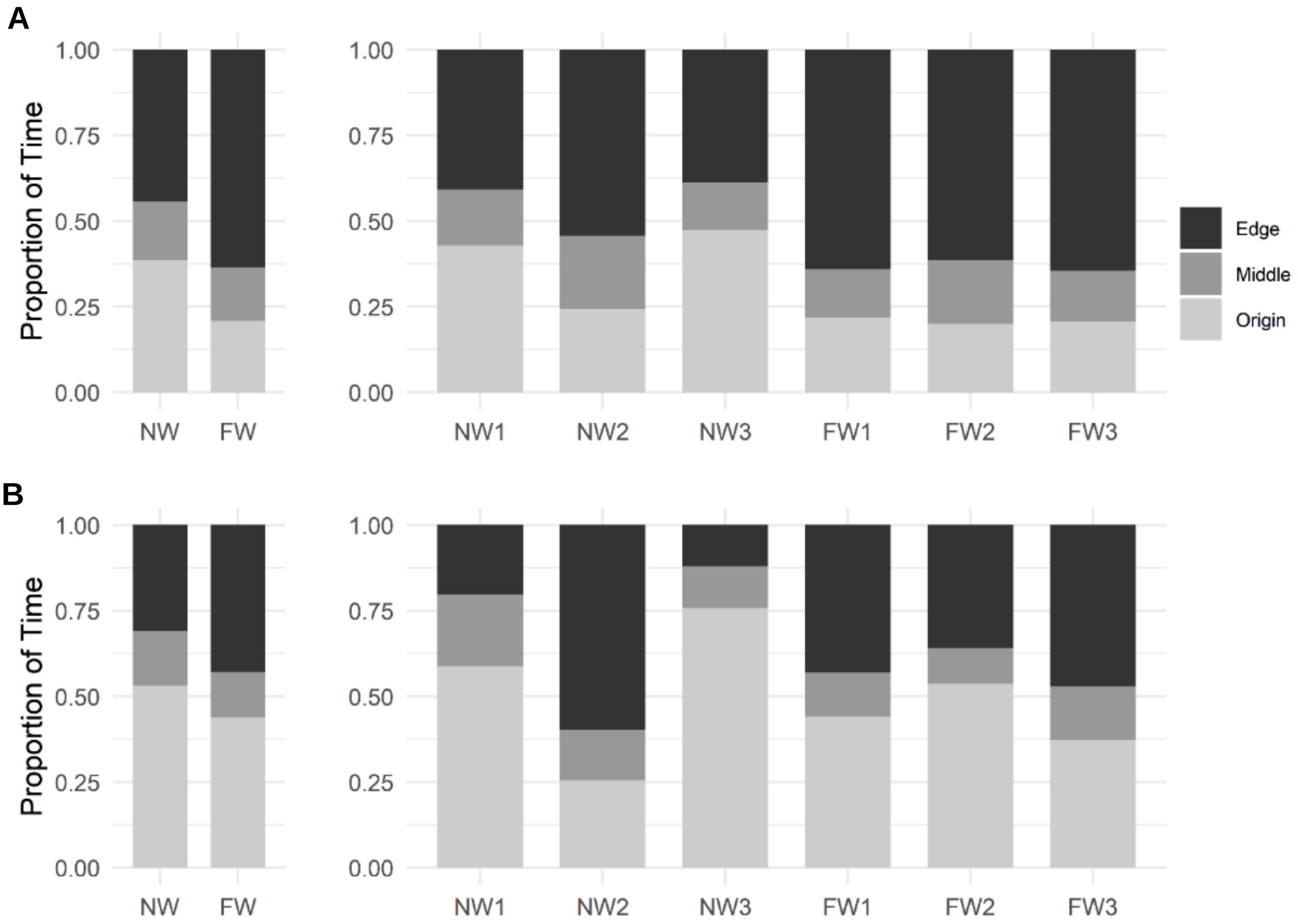

**Figure S6.** The effects of morph genotype on proportions of time spent in each area of the arena by (**A**) 15 day-old and (**B**) 45 day-old juvenile crickets. Plots on the left show data pooled across replicate lines, and line-specific plots are shown on the right.

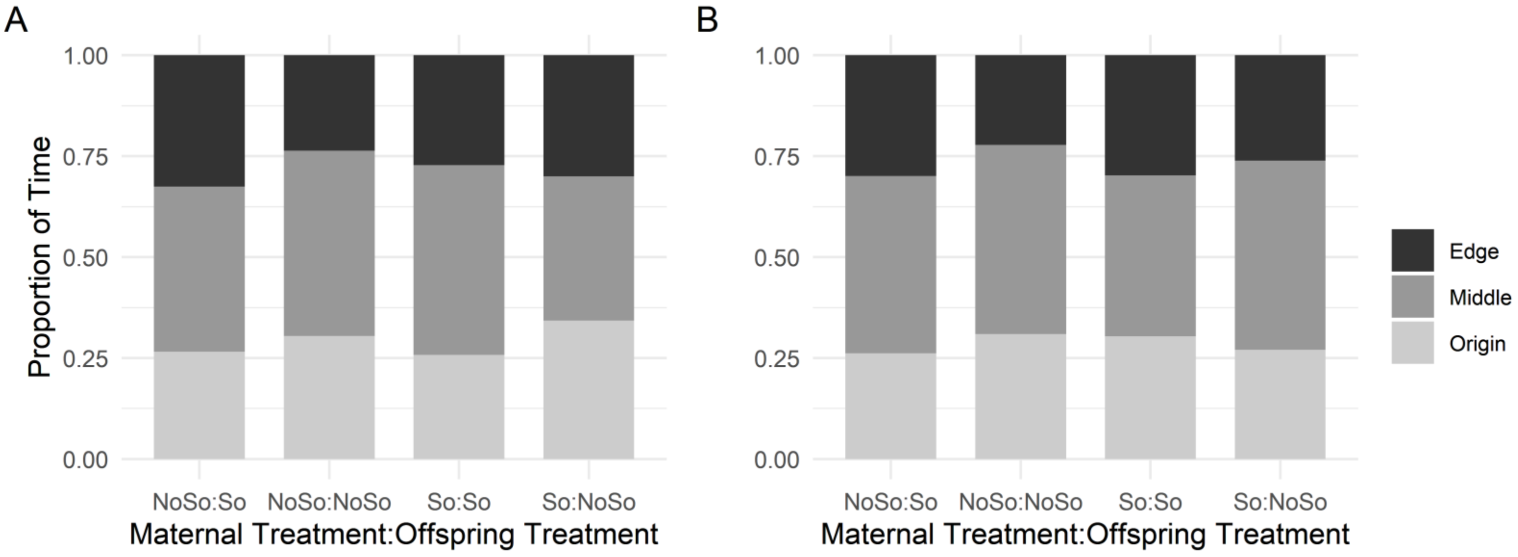

**Figure S7.** The effects of TGP (maternal acoustic treatment) and WGP (offspring acoustic treatment) in (**A**) adult females and (**B**) adult males on proportion of time spent in the three regions of the open field test arena.
